# Conformational switching of Arp5 subunit differentially regulates INO80 chromatin remodeling

**DOI:** 10.1101/2024.05.10.593625

**Authors:** Jeison Garcia, Somnath Paul, Shagun Shukla, Yuan Zhong, Karissa Beauchemin, Blaine Bartholomew

## Abstract

The INO80 chromatin remodeler is a versatile enzyme capable of several functions, including spacing nucleosomes equal distances apart, precise positioning of nucleosomes based on DNA shape/sequence and exchanging histone dimers. Within INO80, the Arp5 subunit plays a central role in INO80 remodeling, evidenced by its interactions with the histone octamer, nucleosomal and extranucleosomal DNA, and its necessity in linking INO80’s ATPase activity to nucleosome movement. Our investigation reveals that the grappler domain of Arp5 interacts with the acidic pocket of nucleosomes through two distinct mechanisms: an arginine anchor or a hydrophobic/acidic patch. These two modes of binding serve distinct functions within INO80 as shown in vivo by mutations in these regions resulting in varying phenotypes and in vitro by diverse effects on nucleosome mobilization. Our findings suggest that the hydrophobic/acidic patch of Arp5 is likely important for dimer exchange by INO80, while the arginine anchor is crucial for mobilizing nucleosomes.

## Introduction

The holocomplex referred to as INO80, with its catalytic subunit known as Ino80, plays a pivotal role in chromatin remodeling important in transcription regulation, DNA repair and replication, nuclear organization and cellular differentiation (1–7). INO80 remodels chromatin by repositioning nucleosomes in a DNA shape and sequence-dependent manner, placing nucleosomes equally separated by the same length of linker DNA and facilitating the exchange of histone dimers (8–13). Deletion of Ino80 hampers ventricular compaction during heart development and correlates with defective coronary vascularization (14). Mutations in YY1AP1, a subunit of INO80, disrupts smooth muscle differentiation and are associated with Grange syndrome, which occasionally manifests as arterial disease (15). Elevated expression of Ino80 is crucial for the proliferation of melanoma (16), small cell lung cancer (SCLC) (17) and colon cancer (18). INO80 increases accessibility at enhancer regions in melanoma and SCLC, thereby facilitating Mediator complex binding (16,17). Additionally, at centromeres INO80 displaces canonical H3 to promote incorporation of CENP-A, and is required for the ectopic incorporation of CENP-A when over overexpressed (19,20). In homologous recombination, INO80 facilitates Rad21 filament formation and DNA resectioning through the removal of H2A.Z (21–23). H2A.Z removal by INO80 also occurs at pericentric regions(24) and certain promoters(25,26), although at many promoters RNA polymerase (RNAP) II primarily displaces H2A.Z (27).

One of the distinctive architectural features of INO80 is the binding of the Arp8 module to linker DNA, which is tethered to the Ino80 catalytic subunit via the HSA domain of Ino80 (28–30). This binding is disrupted by either truncating the N-terminus of Arp8 or by mutating the DNA binding surface of the HSA domain (9,10,31,32). The nucleosome mobilizing activity of INO80 is significantly impaired when the Arp8 module fails to bind linker DNA, suggesting that this mechanism governs INO80’s nucleosome spacing and potentially enables positioning of nucleosomes in a DNA shape-specific manner(8).

Another characteristic architectural element of INO80 is the Arp5 module, which interacts with the histone octamer, nucleosomal DNA at SHL-1, -2 and -3 and extranucleosomal DNA where DNA enters nucleosomes(8,29,33). Arp5 forms a heterodimer with the Ies6 subunit in *Saccharomyces cerevisiae* and contains a flexible grappler domain that exhibits two alternative conformations known as the cross and parallel conformations, as observed by cryo-EM(29). Recently, an improved structure of *Chaetomium thermophilium* INO80:NCP and AlphaFold2 modeling of the grappler domain has provided insights into the closed conformation of the grappler domain. In this model, a region termed the “foot and heel” binds to the acidic pocket of nucleosomes using an arginine anchor (8,34). This model is supported by mutations in the arginine anchor and other regions of Arp5 predicted to interact with nucleosomal DNA effectively blocking nucleosome mobilization by INO80 (34).

The acidic pocket represents a crucial binding site on the nucleosome, characterized by a negatively charged depression. It serves as binding site for various chromatin-associated factors, including viral proteins (35), histone modifiers (36,37) and ATP-dependent chromatin remodelers (29,38–41). Interactions at this site play pivotal roles in regulating transcription, DNA repair and other genomic processes (41–44).

In this study, we delve into the interactions of *Saccharomyces cerevisiae* (Sc)Arp5’s grappler domain by mapping its contacts through site-directed crosslinking from two positions on the histone octamer, one of which lies in close proximity to the acidic pocket. Surprisingly, our investigation reveals the existence of two regions of Arp5 that interact with the acidic pocket that are spatially far apart. These findings suggest the presence of two alternative conformation states of the grappler domain that do not simultaneously bind and sheds light on the dynamic nature of this crucial interaction.

To gain deeper insights, we conducted further experiments to elucidate the functional properties associated with these two conformations, both in vitro and in vivo. Our results provide compelling evidence supporting the notion that these distinct contacts play pivotal roles in modulating the different enzymatic activities of INO80, unraveling the intricate interplay between Arp5’s grappler domain and the acidic pocket of the nucleosome.

## Materials and Methods

### Yeast Strains and Anchor away

Strains used in this study are W303 *(MATα his3-11, 15 leu2-3,112 trp1Δ ura3-1 ade2-1 can1-100) or BY4741 (MATα his3Δ1 leu2Δ0 met15Δ0)* or their derivatives (Supplementary Table S1). Briefly*, Saccharomyces cerevisiae BY4741 (MATα his3Δ1 leu2Δ0 met15Δ0)* strain with a 2 - FLAG (DYKDDDDK) tag epitope attached at the C-terminus of the Ino80 subunit was used for INO80 complexes purification. For Arp5 mutant complexes, Arp5 was replaced in the parental strain by a Kanamycin cassette and the resultant strain was transformed with pRS-416 plasmids using lithium acetate transformation (Supplementary Table S1).

An anchor-away system was employed for the phenotypic characterization of Arp5 mutations (45). Briefly, *S.c. W303 (MATα his3-11, 15 leu2-3,112 trp1Δ ura3-1 ade2-1 can1-100)* strain was modified to tag Arp5 at C terminal with the FRB domain (Supplementary Table S1), using a standard Lithium acetate transformation yeast transformation protocol with a pFA6a-FRB-KanMX6 plasmid (Supplementary Table S1). Transformants were confirmed by PCR. The Arp5-FRB strain was transformed with pRS-416 plasmids expressing varied Arp5 mutants (46).

### Purification of INO80 complexes

Native *Saccharomyces cerevisiae* wild-type and mutant Arp5 complexes were purified using immunoaffinity purification with 2 - FLAG (DYKDDDDK) epitopes attached at the C-terminus of the catalytic subunit Ino80. Yeast strains (Supplementary Table S1) were grown in 100 mL YPD (1 % yeast extract, 2 % peptone, 2 % dextrose, 40 ppm antifoam A and 0.05 % adenine sulfate) for wild type Arp5 or in synthetic complete media-Ura for mutant Arp5-containing complexes. The 100 ml cultures were added to 15L YPD cultures (1 % yeast extract, 2 % peptone, 2 % dextrose, 40 ppm antifoam A and 0.05 % adenine sulfate) and incubated with stirring until the OD600 reached 5-6. Cells were harvested by centrifugation at 4°C, washed with sterile water and H-0.3 buffer (300 mM NaCl, 25 mM Na-Hepes pH 7.8, 0.5 mM EGTA, 0.1 mM EDTA, 2 mM MgCl2, 20 % glycerol, and 0.02 % NP-40) plus protease inhibitors (1 mM PMSF, 1 mM β- mercaptoethanol, 0.5 Na-metabisulphite, 2 μM pepstatin, 0.6 μM leupeptin, 2 mM benzamidine, and 2 μg/ml chymostatin). Yeast spaghetti was prepared from pellets by extrusion into liquid nitrogen and cryogenically ground using a Spex freezer miller 6870. The yeast powder was resuspended with H-0.3 buffer plus inhibitors, and the nuclear proteins were extracted by ultracentrifugation at 100,000 g for 1 h. The supernatant was incubated overnight at 4°C with Anti-FLAG M2 agarose beads (Sigma Aldrich) (10 μl beads per ml extract) in buffer H-0.3 with protease inhibitors. The resin was washed several times with H-0.5 buffer followed by H-0.1 buffer (same composition as H-0.3 buffer but containing 500 mM and 100 mM of NaCl respectively) with protease inhibitors. FLAG-tagged protein complexes were eluted with a 1 mg/mL solution of 3X-FLAG peptide in buffer H-0.1+ 1mM PMSF. The integrity and quantification of the purified complexes was evaluated by 4–20 % gradient SDS–PAGE and SYPRO Ruby staining.

### Western blots

Purified INO80 with wild type Arp5 and Arp5 tagged with HA are resolved on a 20 cm × 20 cm 4–20% Tris-glycine SDS-polyacrylamide gels and transferred onto PVDF membranes using Bio-Rad Trans-Blot® electrophoretic transfer cell for 3 hours at 4 °C, using a transfer buffer containing 25mM Tris, 192mM glycine, 20% methanol, 0.1% SDS (pH 8.3) at 50 V constant voltage. The membranes were blocked with 5% fat-free milk in TBST (20mM Tris-HCl pH 7.5, 150mM NaCl, 0.1% Tween-20), overnight at 4 °C, washed with TBST, and incubated with Anti-HA antibody (Supplementary Table S1) diluted 1:1000 for 1 hour at room temperature. The blots were washed with TBST and developed with SuperSignal™ West Femto Maximum Sensitivity Substrate (Thermo Fisher) before visualization using an Image Quant LAS 4000 (GE healthcare Life Sciences).

### Phenotypic screening

WT and mutant Arp5 cells were grown overnight in 5 ml synthetic complete -Ura media and the OD600 adjusted to 1 by the addition of fresh media. One hundred μl of each culture was taken out and 5-10-fold serial dilutions were made. Next, 1 μL of the original culture (at OD600 = 1) and each of the serial dilutions were aliquoted onto -Ura, HydroxyUracil (HU),6-azauracyl (6-AU) or methyl methanesulfonate (MMS) plates, in presence or absence of rapamycin (8 μg/ml in DMSO). Samples were allowed to dry and then plates were then turned over and left in 30 °C for (37°C for heat shock treatment experiments) 2–3 days until appearance of yeast colonies.

### Peptide mapping of Ino80 subunit with ArgC protease

After label transfer, photoaffinity-labeled Arp5 complexes were denatured with 0.4% SDS and heating at 90 °C for 3 minutes, followed by buffer exchange using Amicon Ultra filters to remove SDS and HA peptides. C-terminal HA-tagged Arp5 was purified by immunoaffinity chromatography. Isolated proteins were washed and resuspended in ArgC incubation buffer containing 50mM Tris-HCl (pH 7.8), 5mM CaCl_2_ and 2mM EDTA. Protein cleavage was initiated by the addition 5mM DTT (final concentration) and varying concentrations of ArgC protease (Promega, sequencing grade) with incubation at 37 °C for 2 hours. Reactions were stopped by the addition of 1 mM PMSF and 10mM EDTA. Immobilized C-terminal fragments were separated from the released fragments and washed three times in the same buffer as the digestion. The bead fractions were resuspended in SDS sample buffer, resolved on 4–20% Tris glycine SDS-polyacrylamide gels, and analyzed by both phosphorimaging, as well as transfer and anti-HA immunoblotting. Apparent molecular masses of the Arp5-HA fragments were estimated by comparing their migration relative to the [^35^S]-labeled Arp5-HA described before (32). Data images in figures are representative of n=3 experiments.

### Binding Assay

Cy5 labeled DNA template was amplified for 70N5 601 DNA templates and nucleosomes were reconstituted. An increasing concentration of INO80 complex (WT or mutant) was incubated with 25 nM nucleosome at 30°C for 30 minutes, in 10 mM Na-HEPES (pH 7.8), 4 mM MgCl_2_, 60 mM NaCl, 0.2 mM EGTA, 0.04 mM EDTA and 8% glycerol. Reactions were analyzed by resolving enzyme-bound nucleosomes from free nucleosomes on 4% native polyacrylamide gels in 1X Tris-EDTA buffer. Data images in figures are representative of n=3 experiments.

### Nucleosome reconstitution and remodeling

Initially 5’ end of the forward primer for 70N5 601 DNA template was labeled with 50μCi of [^32^P]-ATP using T4 Phosphokinase. Then [^32^P]-labeled primer and unlabeled reverse primer was used to amplify the 70N5 601 template DNA. Mononucleosomes were reconstituted with 1.7 μg of 5mg/mL Salmon Sperm DNA and 100 fmol of [^32^P]-labeled 70N5 601 DNA at 37°C with 3-5 μg of recombinant *Xenopus laevis* histone octamers by rapid salt-dilution from 2 M to 280 mM NaCl in 10 minutes steps. The reconstitution was evaluated by native 4% polyacrylamide gel (35.36 acrylamide: 1 bisacrylamide) electrophoresis, followed by phosphorimaging (Typhoon FLA 9500 laser scanner, GE Healthcare Life Sciences). In the case of nucleosome remodeling, the reconstituted nucleosomes (24 nM) were incubated with saturating amounts of INO80 complexes (25 nM) at 30°C for 30 min. Nucleosome sliding was initiated by adding ATP to a final concentration of 80 μM, and remodeling reactions were stopped at the indicated time points by adding 2 mmol of EDTA and 1 μg of salmon sperm DNA (stop mix). Samples were analyzed on 5 % native polyacrylamide gels in 0.5X TBE buffer and phosphorimaged as before.

### ATPase assay

ATPase assays were performed under conditions identical to the remodeling time course assays. After pre-binding, initially a premix of ATP was prepared containing 1mM of cold ATP supplemented with 0.5uL of 3000Ci/mmol γ-^32^P-labeled ATP. ATP mix was added to nucleosome bound complex such that the final concentration of cold ATP was 80μM. Reactions were allowed to proceed for the indicated timepoints. Reactions were stopped by addition of EDTA and SDS to final concentrations of 100 mM and 2 %, respectively. Reactions were spotted onto a polyethyleneimine cellulose plate (J.T. Baker) and developed with 0.5 M LiCl and 0.5 M formic acid.

### Histone Crosslinking

Recombinant *Xenopus laevis* histones with amino acids replaced by cysteine at specific positions (Fig. S1A) were expressed, purified, and reconstituted into octamers with other histones as discussed above. Homogeneous 70N5 WT nucleosomes were reconstituted using the respective 601-nucleosome positioning sequence. The modified nucleosomes were then conjugated with PEAS [N-((2-pyridyldithio)ethyl)-4- azidosalicylamide] (Thermo Fisher), iodinated with [^125^I] and bound to saturating concentrations of WT or mutant Arp5 complexes. Samples were crosslinked by UV irradiation for 3 minutes at 310 nm, 2.65 mW/cm^2^. After crosslinking, the [^125^I]-radiolabel was transferred to the crosslinked proteins by incubating with 100 mM DTT for 30 min at 37°C. Proteins were separated on 4-20% SDS-polyacrylamide gels, dried and viewed by phosphorimaging.

### Peptide pull-down

Biotin-labeled peptides (Supplementary Table S1) were obtained from CPC scientific Inc (San Jose, CA). Streptavidin beads (25 μL) were washed in buffer containing 1X PBS pH 7.4 and 0.01% BSA and incubated with 10 μg of biotin-labeled peptide for 30 min at room temperature with gentle rotation, followed by magnetic pulldown for 2-3 min. The supernatant was discarded and the coated beads were washed 4 times with PBS containing 0.1% BSA and re-suspended at 25 μL final volume. Coated beads were incubated with 5 μg histone octamer in 500μL binding buffer (150mM NaCl, 50mM Tris-HCl, 10% Glycerol, 0.1% NP40) overnight at 4°C with gentle rotation. Next, beads were washed 4 times with 1mL binding buffer on magnetic rack, followed by the addition of 30 mL of 2x Laemilli buffer and boiling for 5 minutes. Samples were analyzed by 15% SDS-PAGE and histones were visualized by using GelCode Blue Stain Reagent (Cat#24590). For antibody competition, histone octamer was pre-incubated with the H2A acidic patch antibody (Millipore 07-146, 1:500 dilution) and Rabbit IgG control for 2 hours before the addition to the peptide coupled beads.

### Peptide competition experiments

WT and mutant LLD peptides were incubated at varied concentrations with reconstituted 601 nucleosomes for 20 minutes at 30°C. INO80 was added to nucleosomes pre-incubated with peptides for 30 min at the same temperature and remodeling initiated by the addition of ATP to a final concentration of 80 μM. After 10 min, reaction was stopped as described previously and resolved on a 5% native polyacrylamide gel in 0.5X TBE buffer, followed by phosphorimaging.

### Site-directed mapping at H2B53–DNA contacts

Histone octamers with cysteine at residue 53 of H2B were conjugated to p-azido phenacyl bromide (APB) immediately after octamer refolding. Reactions contained 80 nM wild type or mutant Arp5 complexes and 80nM nucleosomes and were incubated at 30 °C for 30 min in conditions. Nucleosome movement was initiated by adding 80 μM ATP and stopped with 300 ng/μl sonicated salmon sperm DNA and 10 mM EDTA at the indicated time points. For site-directed histone-DNA crosslinking, samples were irradiated with UV for 3 min (310 nm, 2.65mW cm−2). Samples were denatured with 0.1% SDS at 37 °C for 20 min in 30 mM NaCl and 20 mM Tris-HCl (pH 8.0). Crosslinked protein-DNA were enriched and separated from un-crosslinked DNA by phenol-chloroform (4:1) extraction. The aqueous phase containing un-crosslinked DNA was discarded. Crosslinked DNA was ethanol precipitated with 1M LiCl in the presence of sheared salmon sperm DNA as carrier. Crosslinked DNA was cleaved with 1 M pyrrolidine (Sigma) at 90 °C for 15 min. DNA samples were analyzed alongside a sequence ladder made from the same DNA on a denaturing 6.5% polyacrylamide gels containing 8 M urea. Gels were visualized by phosphorimaging and quantified using ImageQuant software (Version 5.2). Data images in figures are representative of n=3 experiments. %Net loss of signal for WT or mutants on the entry side was calculated using formula 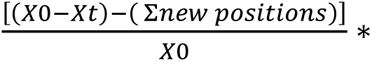 100 where X0 is the crosslinking at time 0 s and Xt is the crosslinking at time ts where t= 0 to 2560s.

### Alpha2 Fold Modeling of *Saccharomyces cerevisae* (Sc) Arp5

The Alpha2 Fold predicted structure for Sc Arp5 comes from the AlphaFold Structure Database of EMBL-EBI and is AF-P53946-F1 model v4_1. This structure was last updated in AlphaFold DB version 2022-11-01 and created with the AlphaFold Monomer v2.0 pipeline (47). The confidence of the structure is shown using the predicted local difference test (pLDDT) and most of the grappler domain has pLDDT scores ranging from 70 to 90 in the high confidence range (Figure S6). The AlphaFold2 structure of Sc Arp5 is aligned using Pymol 2.5.4 from Schrodinger, LLC to the Ct Arp5 structure that was obtained using a combination of cryo-EM and AlphaFold2 modeling to confirm the overall similarity between the two structures (Figure S6)(9).

## Results

### Three distinct parts of the grappler domain interact with two regions of the histone octamer

We probe the interactions of the Arp5 subunit with the acid pocket of nucleosome and adjoining area by site-specific histone to INO80 crosslinking. Photoreactive nucleosomes are made by first constructing seven different histone octamers, changing the native Cys 110 of histone H3 to Ala and changing another histone amino acid residue located at an exposed region of nucleosomes to Cys (Figure S1A). An ^125^I labeled photoreactive reporter is covalently linked to Cys and after crosslinking the radiolabel is transferred to the crosslinked INO80 subunit(s). The INO80 subunit most widely crosslinked at these positions is Arp5 and is most efficiently crosslinked to residue 89 and 80 of respectively histones H2A and H3 (Figures 1B and S1B). Residue 89 of histone H2A (H2A 89) is located on the edge of the acidic pocket of nucleosomes and residue 80 of histone H3 (H3 80) is adjacent to the region where the DNA binding domain of Arp5 binds to Super Helical Location (SHL)-3 and -2, consistent with prior cryo-EM data of INO80-nucleosome complexes(8,10,29,33,34).

**Figure 1.**
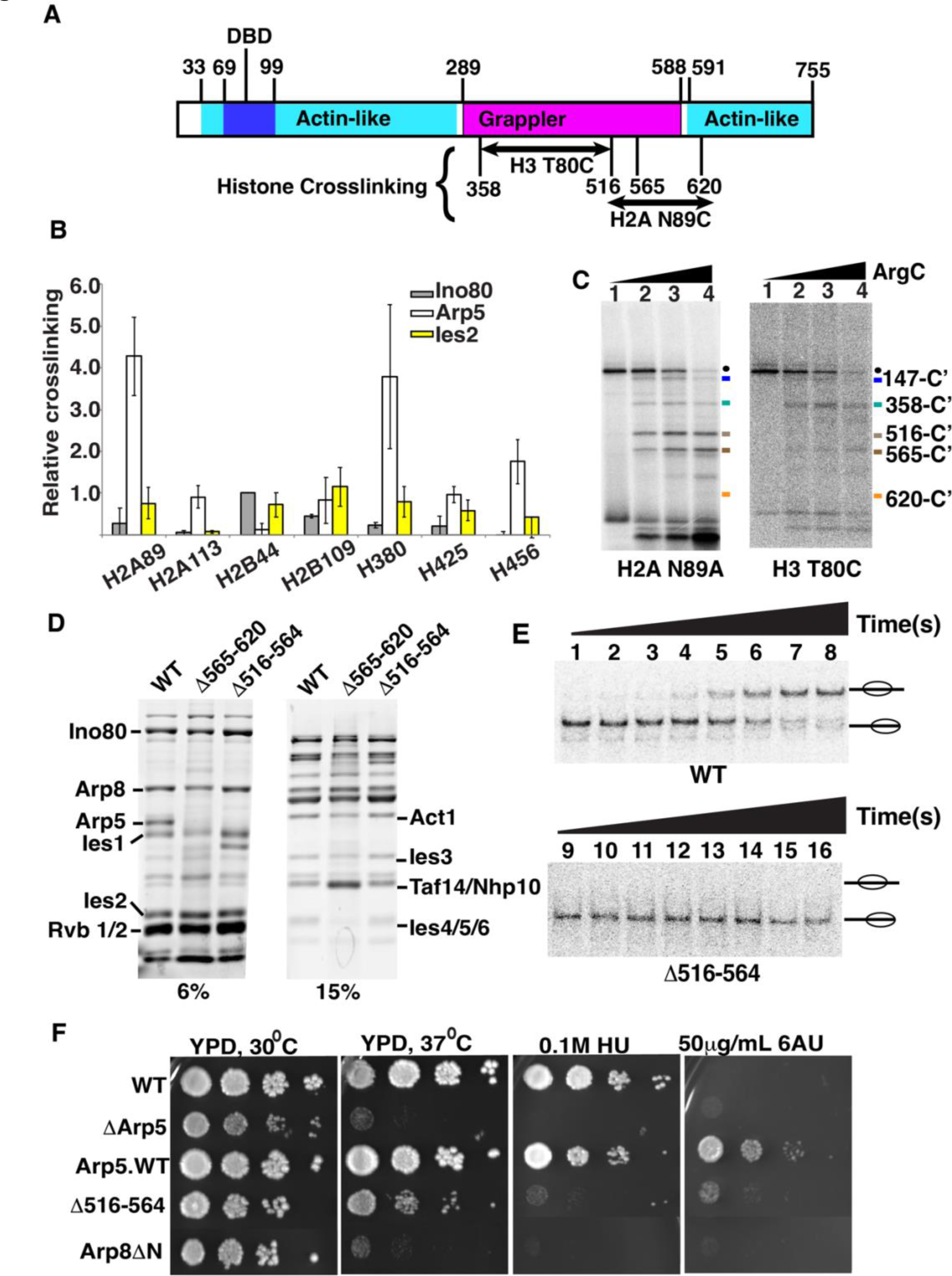
Two regions of Arp5’s grappler domain associate with the acid pocket of nucleosomes. (A). The domain organization is shown for the Arp5 subunit of INO80 with the actin-like portion highlighted in cyan and the grappler domain in purple. The DNA binding domain of Arp5 is indicate (dark blue. The Arp5 regions crosslinked to residue 80 of histone H3 and residue 89 of H2A are labeled. (B) The interactions of Arp5 with the histone octamer face of nucleosomes are probed by site-specific histone crosslinking. The relative efficiency of crosslinking Arp5, Ies2 and Ies6 is shown for three replicates. Error bars represent the mean ± SD.(C) The regions of Arp5 crosslinked to histones are mapped by immobilizing the C-terminus of Arp5 and partial digesting with Arg-C. Labeled proteolytic fragments of Arp5 are separated on a 4-20% SDS-PAGE and visualized by phosphorimaging. (D) The two regions are individually deleted, and the complex purified using by FLAG affinity chromatography. (E) Nucleosome mobilizing activity of the INO80 complex with Arp5 ^Δ516-564^ (lanes 1-8) is compared to wild type INO80 (lanes 9-16) using EMSA. (24nM) of nucleosomes were saturated with 25nM of INO80 complex (WT or mutant) with addition of 80 μM ATP. (F) The phenotype of the yeast strain with Arp5 ^Δ516-564^ was compared to wild type Arp5 (WT, Arp5 WT), N-terminal truncation of Arp8 (Arp8 ΔN) and the absence of Arp5 (ΔArp5).

Next, we determine the region of Arp5 contacting nucleosomes at these two primary crosslinking sites by peptide mapping. A hemagglutinin (HA) epitope is fused onto the C-terminus of Arp5 for recovering only the C-terminal fragments of Arp5 after cleavage with Arg-C specific protease, which did not perturb binding of Arp5 at these two sites (Figures 1C, S1C and S2C). The proteolytic fragments are separated by SDS-PAGE along with a set of Arp5 specific molecular weight markers and visualized by phosphorimaging (Figures 1C and S1D-E). Two different regions of Arp5 are crosslinked to H2A 89 that are amino acid 516 to 564 and 565 to 620, both with limited and extensive proteolytic digestion (Figure 1C, compare lanes 2 and 4). H3 80 crosslinked to amino acids 358 to 516 of Arp5 and all three crosslinked regions are in the grappler domain of Arp5.

The two regions crosslinked near the acidic pocket are analyzed further by deleting each region and checking its effect on INO80 complex integrity and nucleosome mobilizing activity plus the in vivo activity of truncated Arp5 under different Arp5 dependent conditions. INO80 complex integrity is maintained when amino acids 516-564 (Arp5^Δ516-564^) are deleted, but when amino acids 565-620 (Arp5^Δ565-620^) are deleted both Ies6 and truncated Arp5 are lost from the complex (Figure 1D). Previously, deletion of parts of Arp5 in the grappler of Arp5 had been shown to disrupt complex integrity similar to that observed here and we didn’t analyze further the Arp5^Δ565-620^ complex due to its lack of complex integrity (48).

In the Arp5^Δ516-564^ INO80 complex, nucleosome mobilization is uncoupled from ATP hydrolysis as nucleosomes are not mobilized to any detectable extent even after an extended time, whereas the rate of ATP hydrolysis is decreased only 3.8-fold versus wild type INO80 (Figure 1E and S2D-E and Table 1). The region from 516 to 564 is also shown to be required for Arp5 to associate near the acidic pocket as seen by the loss of Arp5 crosslinking to residue 89 of H2A with the Arp5^Δ516-564^ complex (Figures S2A-B).

Wild type Arp5 is required in vivo under heat, transcription elongation or DNA replication stress conditions as shown with growth spot assays of wild type Arp5 and ΔArp5 strains (Figure 1F). The Arp5^Δ516-564^ mutant has similar growth defects to that of complete deletion of Arp5 or truncation of the N-terminal part of Arp8 (Arp8ΔN), consistent with the grappler region from amino acids 516-564 being required for the in vivo activity of Arp5 under all three conditions (Figure 1E).

### An arginine anchor in the grappler domain is required for INO80 mobilizing nucleosomes and for Arp5 contacting the acidic pocket of nucleosomes

We scan for residues that are evolutionarily conserved in Arp5 from amino acid 514 to 565 to guide us in determining the critical residues important for Arp5 activity (Figure 2A). We also focus on arginine residues given their potential to interact with the acidic pocket of nucleosomes including arginines 482, R488 and R496 that are adjacent, which based on structural modeling have been predicted to be the arginine anchor that binds the acidic pocket(33,34). Both arginines 488 and 496 are well conserved up to *D. melanogaster* but not in *M. musculus* or *D. melanogaster* and arginine 482 is similar but is sometime replaced with lysine (Figure 2A). There are also two arginines at 514 and 516 that are conserved only in fungi. Next, we observe a hydrophobic/acidic patch from amino acid 529 to 541 that is conserved up to *D.* melanogaster and another acidic patch that is only conserved in fungi from amino acid 550 to 554. We mutate sets of these amino acids to alanine to find the residues critical for Sc-Arp5’s in vivo activity (Figure 2A). An anchor-away system is used to rapidly remove wild-type Arp5 and acutely expose cells to mutant Arp5. The effectiveness of anchoring away Arp5 is shown with growth spot assays before and after removal of endogenous Arp5 (-/+rapamycin) in the absence of any plasmid and with the empty plasmid (pRS416) and Arp5 plasmid (pRS416 Arp5). The anchor away system for Arp5 works well as shown when complementation is tested under heat stress (37 °C), inositol and hydroxy urea selection conditions (Figure 2B).

**Figure 2.**
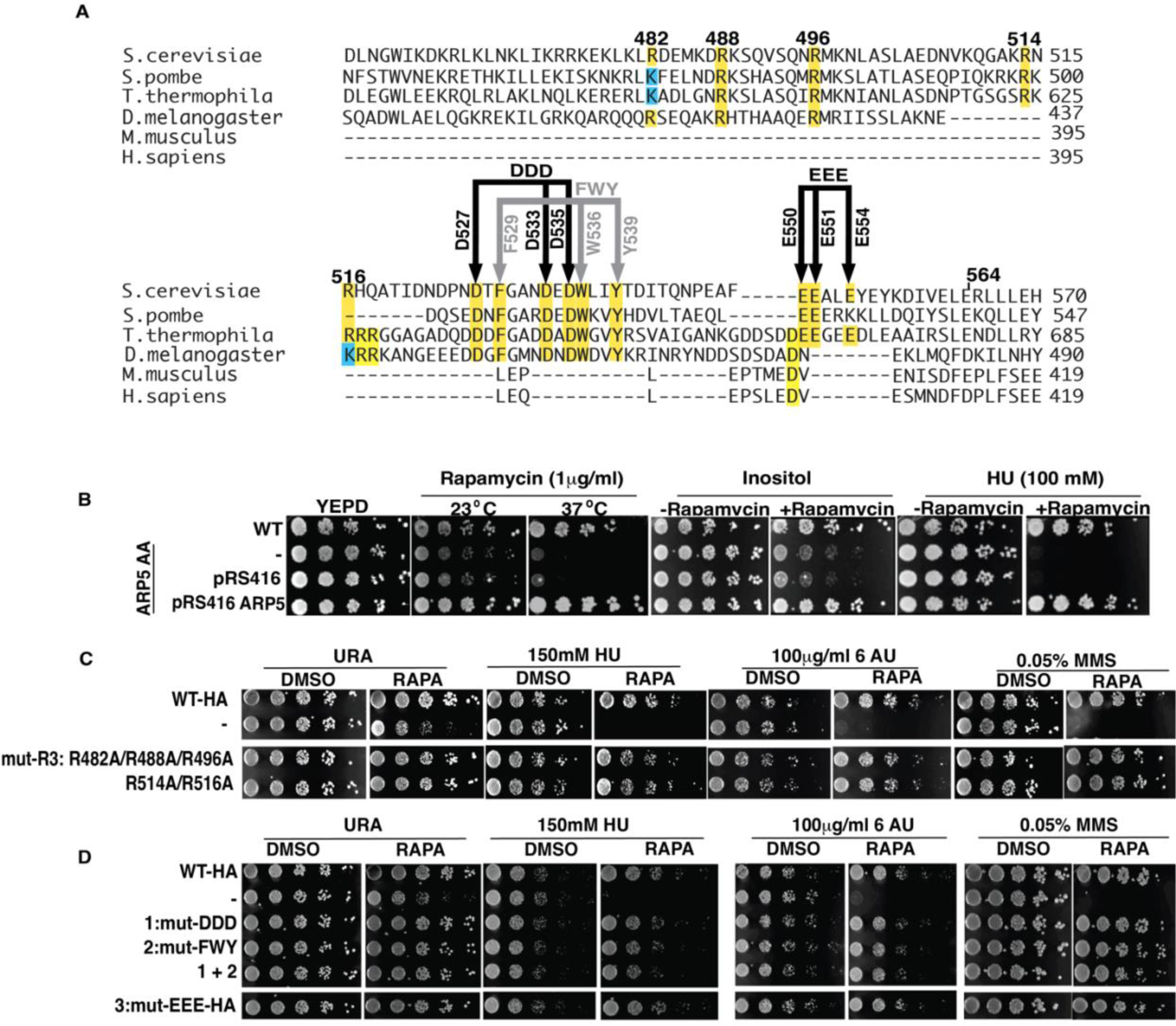
The arginine anchor is not required in vivo for Arp5’s role in DNA replication, DNA damage repair and transcription elongation. (A) Various arginines mutated to alanines in Arp5 are tested in spot assays to determine their impact on the in vivo activity of Arp5. The region spanning amino acids 516-564 has several regions where the sequence is conserved and are probed by mutating to alanine. (B) An Anchor Away (AA) system is used to acutely test cell viability of various mutant Arp5 under different conditions where wild type Arp5 is required for cell proliferation. (C- D) The effect of different mutations in the grappler domain are tested using the AA system with (RAPA) and without (DMSO) added. The abbreviations are as follows: HU-hydroxy urea, 6AU – 6 amino uracil and MMS – methyl methanesulfonate.

We find the putative arginine anchor or both Arg 514 and Arg516 are not required for the in vivo activity of Arp5, which had not been tested previously in vivo for Ct Arp5 (Figure 2C) (34). In the region from amino acid 529 to 541, we mutate hydrophobic amino acids F531, W538 and Y541 and separately the acidic amino acid residues D529, D535, and D537 to alanine and find these residues are also not required for Arp5 function in DNA replication stress, DNA damage repair or transcription elongation either separately or when both groups are combined (Figure 2D). The other moderately conserved acid patch is similarly not required for Arp5 under these same conditions. It is surprising that none of the conserved amino acid residues tested appear to be important for the in vivo activity of Arp5, especially since mutation of the comparable arginine anchor in Ct Arp5 was previously seen to significantly interfere with INO80 remodeling activity(34).

We purified Sc INO80 with R482, R488 and R496 (R3) changed to alanine to probe the functional role of the arginine anchor, which does not affect complex integrity (Figure S3A). The arginine anchor is required for INO80 to mobilize nucleosomes as shown by EMSA as well as to hydrolyze ATP using the mutant complex (Figures 3A-B and S3). The rate of ATP hydrolysis is decreased 4.8-fold by mutating the arginine anchor compared to wild type INO80; whereas it is not possible to do the same comparison for nucleosome mobilization because of how defective the arginine mutant INO80 is compared to wild type (Table 1). Site-directed histone crosslinking to INO80 confirms mutation of the arginine anchor blocks the association of Arp5 near the acidic as seen by reduced crosslinking to residue 89 of histone H2A (Figures 3C and S3C). The loss of Arp5 binding seen with mutation of the arginine anchor is specific to Arp5 as it does not change the extent at which Ies6 or Ies2 is crosslinked to residues 89.

**Figure 3.**
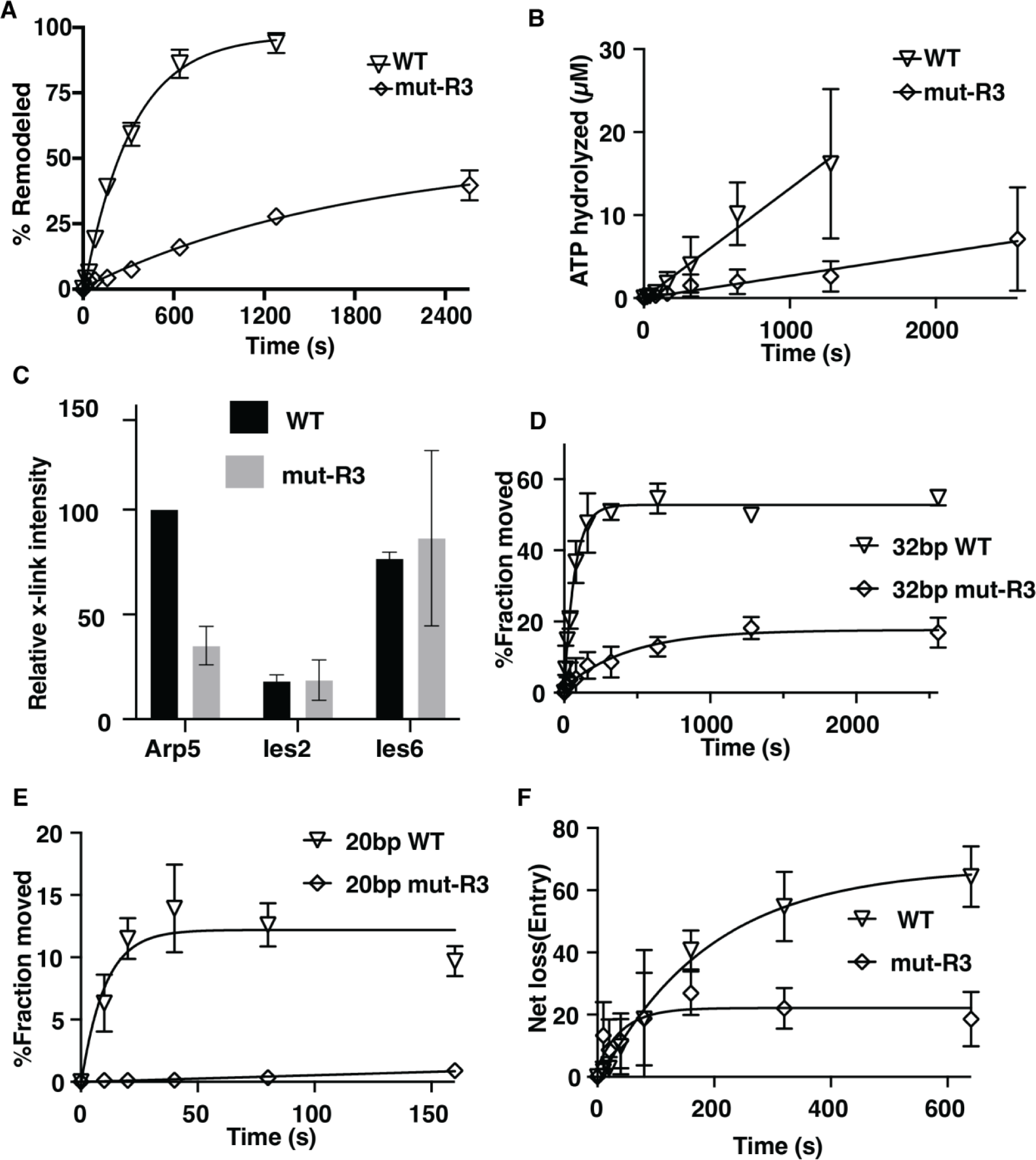
Arginines 482, 488 and 496 of Arp5 are important for Arp5 binding to the acid pocket and for INO80 mobilizing nucleosomes. (A) The graph shows the extent of nucleosomes mobilized by INO80 with wild type (WT) and mutant Arp5 (R3) that has arginine 482, 488 and 496 mutated to alanine using EMSA (see Figure S3B). 50nM of nucleosomes were incubated with 75nM of INO80 complex with addition of 80 μM ATP (B) The ATPase activity of wild type and mutant INO80 was measured using thin layer chromatography under similar condition to that in (A). (C) The interactions of Arp5 with the histone octamer face of nucleosomes are probed for WT and R3 mutant by site-specific histone crosslinking. The relative efficiency of crosslinking Arp5, Ies2 and Ies6 is shown for three replicates. (D-E) The extent of DNA movement on the (D) exit and (E) entry side is plotted versus time using a photoreactive probe attached to residue 53 of histone H2B for wild type and R3 mutant Arp5 containing INO80 complex. (E) The net loss of DNA crosslinking on the entry side is plotted versus time. Three replicates are performed for each experiment and error bars represent the mean ± SD.

The role of the arginine anchor for mobilizing nucleosomes by INO80 is investigated further by tracking DNA movement and displacement inside of nucleosomes during INO80 remodeling using nucleosomes with a photoreactive reporter attached to residue 53 of histone H2B(49,50). The photoreactive reporter detects DNA movement by crosslinking and cleaving DNA at the most proximal nucleotide, which is used to take “snapshots” as DNA translocates pass by this location or is displaced. DNA movement is mapped 53 bp from the dyad axis on both sides where DNA exits and enters nucleosomes during remodeling (Figure S4). We observe DNA movement is strongly repressed at the exit side of nucleosome by the arginine anchor mutant and moves less than 20% of nucleosomes compared to over 50% with a rate that is 6.4-fold slower for an overall reduction of 19-fold (Figure 3D). On the entry side of nucleosomes, the reduction in moving DNA 20 bp is even more dramatic as there is no significant movement detected with the arginine anchor (Figure 3E). DNA is displaced from the histone octamer by INO80 on the entry side as seen by a loss of DNA crosslinking to residue 53 of H2B, which is initially the same for arginine anchor and wild type Arp5 (Figure 3F). The arginine anchor mutant complex however stops after 80 s while with the wild type complex DNA displacement continues to increase. These data suggest the arginine mutant is defective at all stages in nucleosome mobilization that can be detected by site-directed crosslinking.

In summary, we see the arginine anchor of Sc Arp5 is required for nucleosome mobilization by INO80, but nucleosome mobilization does not appear to be the activity of INO80 that accounts for its role in relieving DNA replication stress or assisting in DNA damage repair or potentially transcription elongation.

### A hydrophobic/acidic patch in the grappler domain is required for the in vivo activity of Arp5

We scan for conserved amino acids in the second region that crosslinks to the acidic pocket from amino acid 565 to 620 to find if there are residues that when mutated reflect the phenotype observed by truncation of Arp5. Arginines 618 and 620 are well conserved including in *M.* musculus and *H. sapiens* and when mutated to alanine do not have an Arp5 dependent phenotype (Figures 4A-B). Arginines 565, 596 and 599 are less conserved in this region and when changed to alanine also fail to show an Arp5 dependent phenotype. A short hydrophobic/acidic patch encompassing Leu 567 and 568 and Glu 571 (LLD) when mutated to alanine displays a phenotype consistent with the absence of Arp5 for DNA replication and transcription elongation stress and DNA damage repair (Figure 4A-B). The LLD region appears to behave differently than the arginine anchor based on the phenotypic assays.

**Figure. 4.**
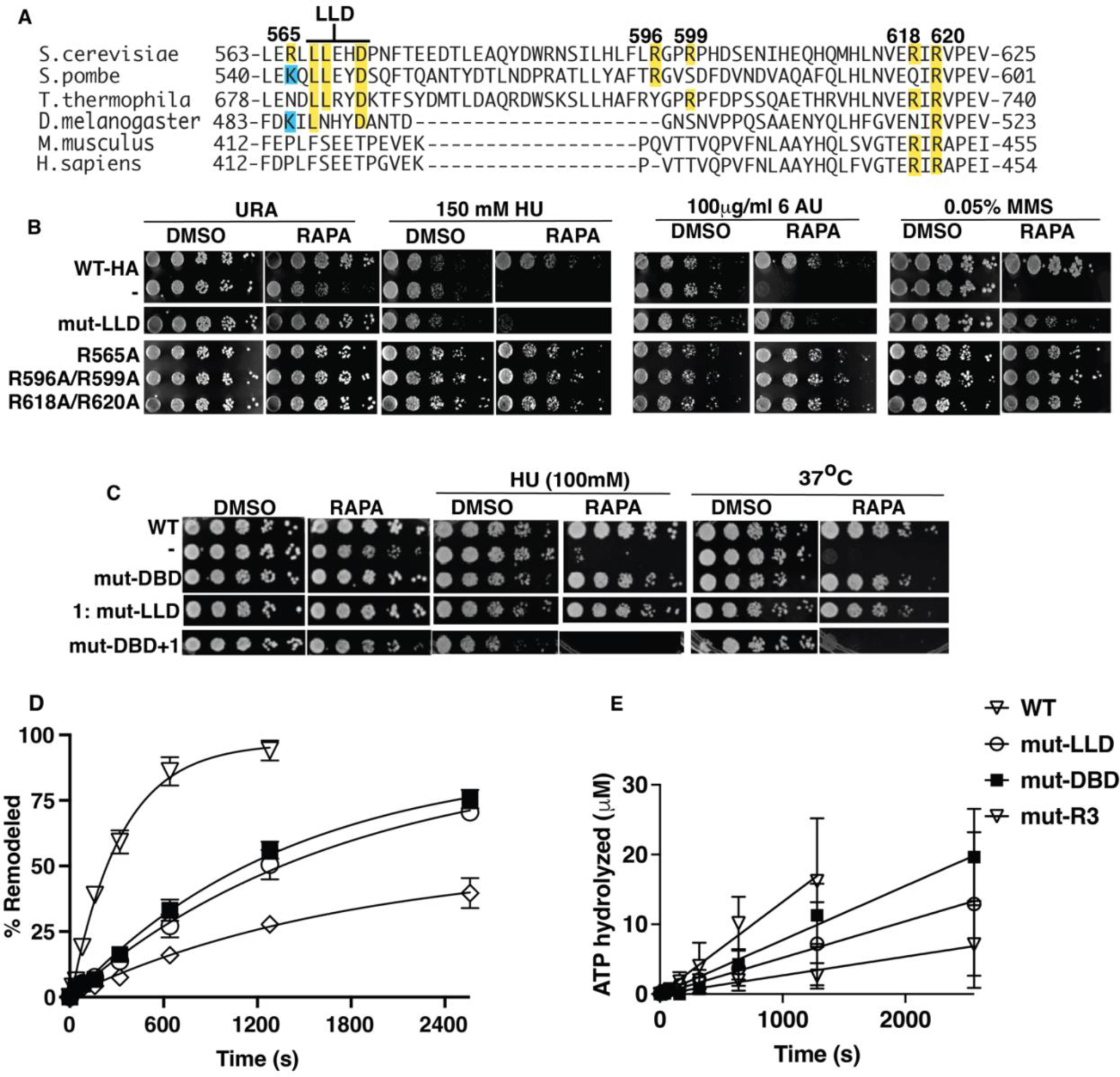
Mutation of Leu 567, Leu 567 and Asp5 to alanine in the grappler domain negatively impacts both the in vivo and in vitro activity of Arp 5. (A) The sequence homology for Arp5 spanning the region from amino acid 565 to 620 of Sc Arp5 with Arp5 from other organisms is shown. *Saccharomyces cerevisiae* (CAA95933.1), *Schizosaccharomyces pombe* (CAB44762.1), *Thermochaetoides thermophila* (XP_006693704.1), *Drosophila Melanogaster* (NP_650684.1), mouse (NP_780628.3) and human (NP_079131.3). (B-C) INO80 with wild type Arp5 is compared to mut LLD with Leu 567, Leu 568 and Asp 571 mutated to Ala, mut DBD with Arg 71, Arg 73, Lys 77, Arg 93 and Arg 97 changed to Ala (DBD) to block it from binding DNA at SHL-2/-3 and mut R3 as described in Figure 3. Two sets conserved arginine residues of (1) 596 and 599 and (2) 618 and Arg 620 are mutated to Ala and screened using spot assays. In (C) The phenotype of LLD and DBD combined are examined. (D) The nucleosome remodeling activity of wild type (WT) and mutant (mut)-LLD or (mut)-DBD INO80 are measured using EMSA and the extent of mobilized nucleosome plotted versus time (see Figure S4C). 50nM of nucleosomes were incubated with 75nM of INO80 complex with addition of 80 μM ATP. (E) The ATPase activity of wild type and mutant INO80 was measured under similar condition to (D). Three replicates are performed for each experiment and error bars represent the mean ± SD.

Mutant LLD containing INO80 complexes are purified and observed to retain complex integrity (Figure S3A). We also purify the DBD Arp5 mutant version of INO80 where binding to nucleosomal DNA at SHL-2/-3 is eliminated by mutating Lys 69, Arg 71, Arg 73, Lys 77, Arg 93 and Arg 97 to Ala, similar to that performed previously for *C. thermophilum* INO80, and compare it to the LLD mutant INO80 complex (29). The affinity of mutant LLD and DBD complexes for nucleosomes is equivalent to that for wild type Arp5 complexes (Figures S5A-B). Mutation of LLD and DBD negatively impacts the nucleosome mobilizing activity of INO80 less than mutation of the arginine anchor of Arp5 and mutant LLD and DBD decreased the mobilizing activity 5.5- and 4.4-fold respectively compared to wild type Arp5 (Figures 4D and S5C and Table 1). The ATPase activities of the DBD, LLD and R3 mutants are decreased from 1.7 to 4.8-fold compared to wild type Arp5 (Figure 4E and Table 1). The differences in nucleosome mobilizing activity between mutant R3 and LLD does not account for mutant LLD having a pronounced phenotype typical of loss of Arp5.

We examine the effects of mutating the LLD in the grappler region and DBD of Arp5 by growth spot assays using a lower HU concentration where the individual sets of mutations do not have a noticeable effect (Figure 4C). The conditional synthetic lethality seen by combining the LLD and DBD mutations suggest these two parts of Arp5 regulate the in vivo activity of Arp5 in distinct ways. A similar synthetic lethality is also observed for heat shock sensitivity when the DBD and LLD mutations are combined (Figure 4C).

### Evidence for alternative conformations of the grappler domain of Arp5 with distinct corresponding activities in vivo

To better understand the LLD region of Arp5, we model the structure of *Saccharomyces cerevisiae* (Sc) Arp5 using AlphaFold2 and align it with the structure of Arp5 from *Chaetomium thermophilum* (Ct) (Figures 5A-B and S6A-B). The Ct Arp5 structure is based on both the cryo-EM structure of Ct-INO80 bound to nucleosomes coupled with AlphaFold2 modeling and displays what is referred to as the cross configuration of the grappler domain (9). In the cross grappler conformation of Arp5, the LLD region is well removed from the acidic pocket of nucleosomes and the adjoining residue 89 of H2A (Figures 5C). The arginine anchor comprising Arg 482, Arg 488 and R496 of Arp5 binds to the acidic pocket when the grappler is in the cross conformation (Figure 5C). The spatial distances however between the arginine anchor and LLD suggest these cannot both simultaneously bind to the acid pocket of nucleosomes and might instead be representative of two alternate configurations of the grappler domain of Arp5.

**Figure 5.**
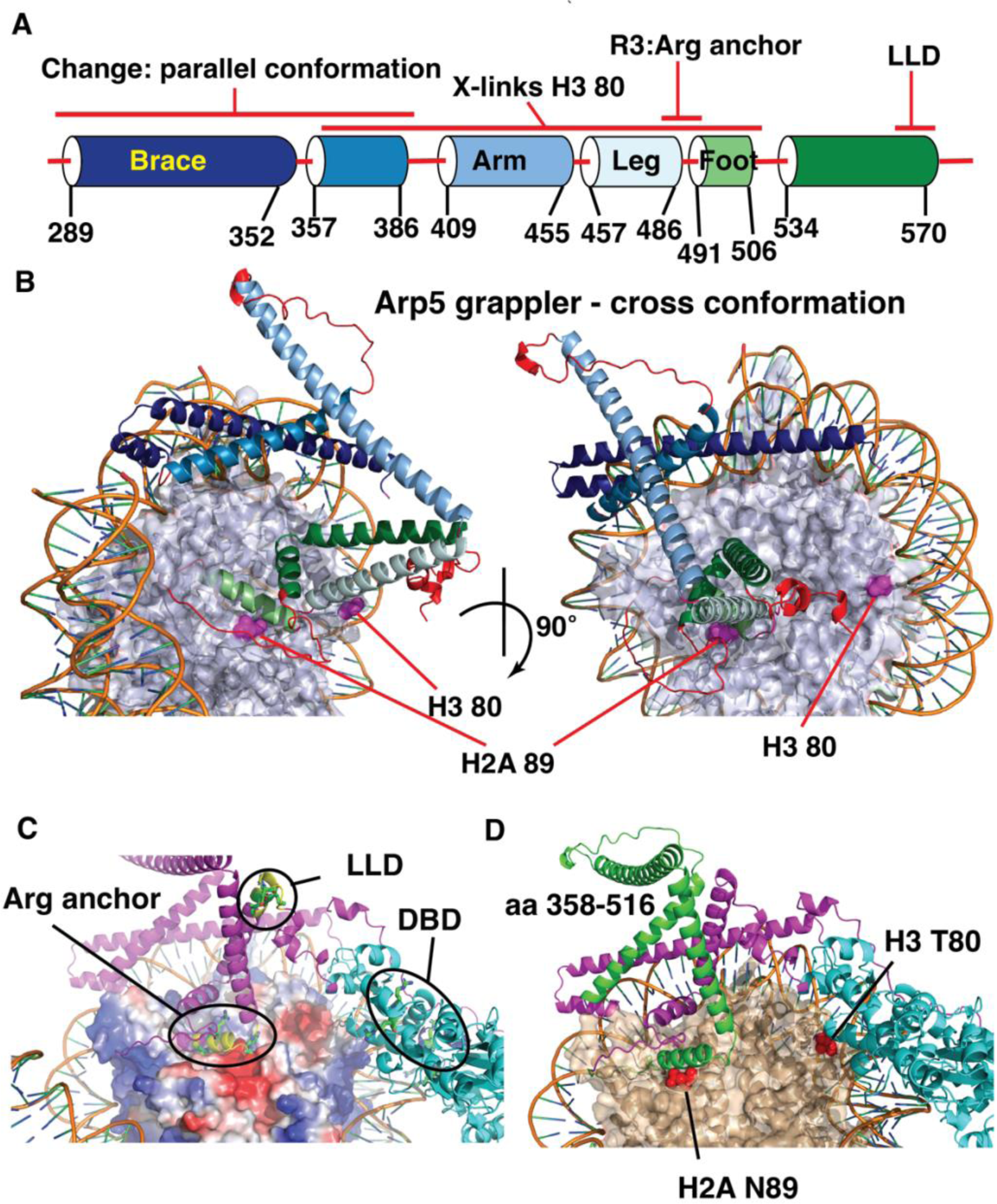
The LLD and arginine anchor (R3) are spatially separated in the grappler domain of Arp5. (A) Schematic shows the alpha helices and loops that compromise the grappler domain of Arp5 based on the closed conformation. Names for different helices are shown along with the positions of the arginine anchor (R3) and LLD region as well as the region crosslinked to H2A 89 and H3 80. (B) The structure of the crossed conformation of the grappler domain of Arp5 shown and the color coding corresponds to that in (A). The location of residue 80 of H3 and 89 of H2a are highlighted. (C) The location of the arginine anchor, DNA binding domain (DBD) and the LLD region are indicated. (D) The same orientation of the grappler domain in (C) is shown with the region crosslinked to residue 80 of H3 highlighted in green and the rest of the grappler domain in purple and the actin-like portion of Arp5 in cyan.

The region of Arp5 crosslinking to residue 80 of H3 indicates the grappler domain acquires a conformation that diverges significantly from the cross conformation. The region spanning amino acids 358 to 515 crosslinked to H3 80 in the cross conformation is close to residue 89 of histone H2B and not residue 80 of H3 (Figure 5D). Next, we want to determine if the LLD region binds to the acidic pocket as might be indicated by the region of Arp5 crosslinked to H2A 89 and if this interaction is part of the alternative grappler conformation observed by Arp5 crosslinking to H3 80.

The model of Sc Arp5 also reveals Arg514 and Arg514 may interact with nucleosomal at the dyad and exit sites of nucleosomes and Arg618 and Arg620 are in surface exposed region of the DBD (Figure S7B-C). Arg 301, Arg 307 and Arg 316 of the grappler domain binds to nucleosome on the entry side of nucleosomes and are similar to a region seen in the cross conformation of Ct Arp5 (Figure S6B). Previously with Ct Arp5, these arginines were found to be required for the nucleosome mobilizing activity of INO80(34). We examined the potential importance of these arginines for Arp5 activity in vivo by mutating them to alanine and using the Anchor-Away system as before. These three arginines are found not to be important for Arp5 activity involved in DNA replication stress response, transcription elongation and DNA repair (Figure S7E).

### The hydrophobic/acid patch of Leu567, Leu568 and Glu571 (LLD) of Arp5 interacts with the acidic patch of nucleosomes

Mutations in LLD disrupt crosslinking of Arp5 to H2A 89, consistent with the LLD directly associating with the edge of the acidic pocket, while not interfering significantly with Arp5, Ies2 or Ies6 crosslinking to residue 80 of histone H3 (Figures 6A-B and S8A- B). The DBD mutant does not alter crosslinking at either histone site as expected based on these mutations targeting specifically Arp5 binding to nucleosomal DNA. DBD and LLD mutants together have a synergistic effect by disrupting Arp5 crosslinking at the H3 80 site, indicating the combination of Arp5 binding to nucleosomal DNA and LLD potentially binding to the acidic pocket contributes to the interactions of Arp5 near residue 80 of histone H3 (Figures 6A-B). These data are consistent with Arp5 interactions near residue 80 being part of the same alternative conformation of the grappler domain as the LLD domain interacting with the acidic pocket. The more extensive loss of Arp5 nucleosomal interactions when combining both the LLD and DBD mutations may explain why when combined these two mutants have a conditional synthetic lethality phenotype (Figure 4B).

**Figure. 6.**
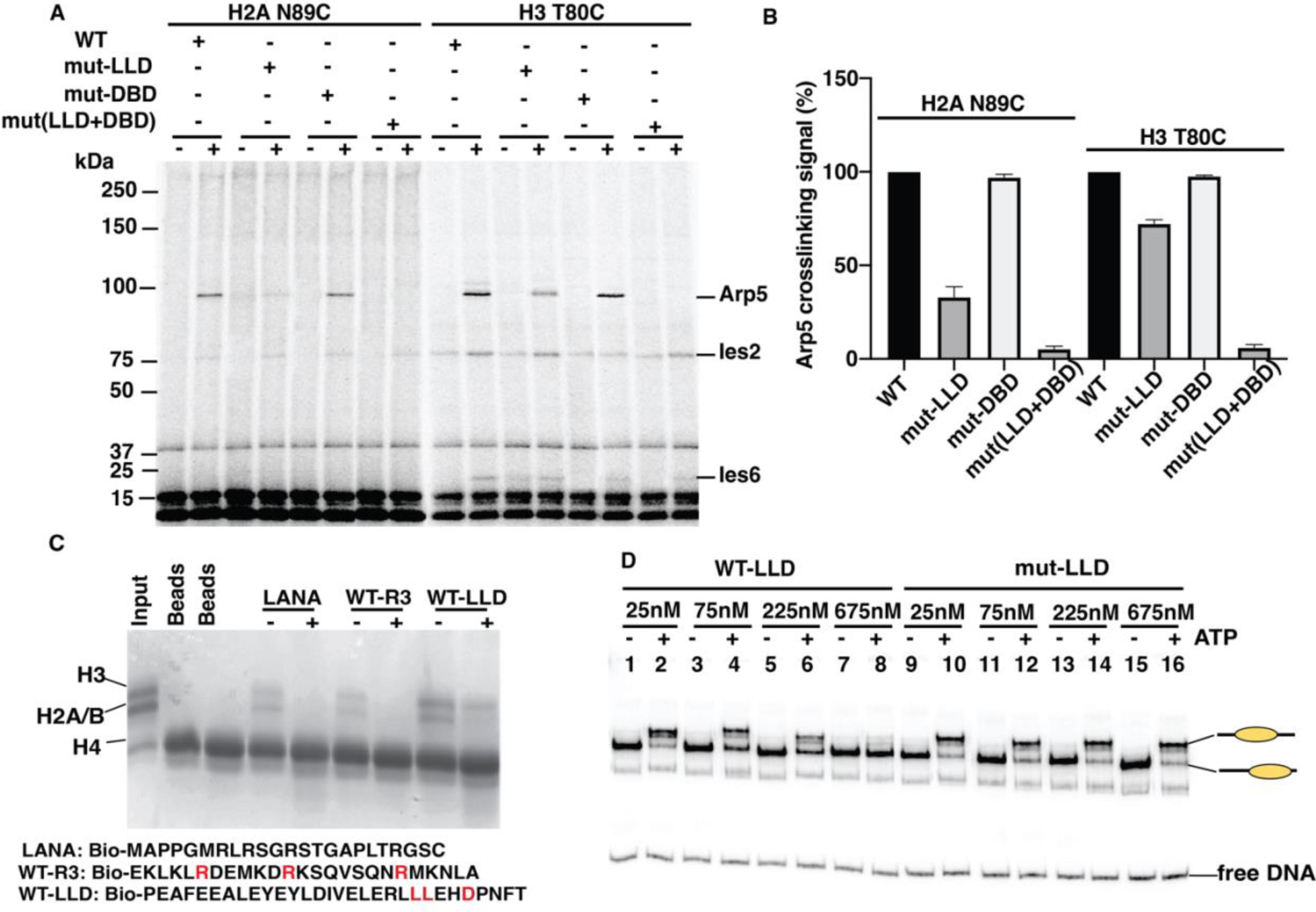
The LLD part of the Arp5 grappler domain is required for Arp5 binding to the acidic pocket of nucleosomes and can independently bind histones. (A-B) Binding of Arp5 near the acid pocket (H2A N89C) and another lateral position on the histone octamer (H3 T80C) is probed by site-specific histone crosslinking for wild type (WT) and mutant LLD and DBD (mut-LLD and mut-DBD). (C) Pulldown assays are done with histone octamer (5μg) and one of three biotinylated peptides; LANA and two others encompassing the region of the Arg anchor and the LLD patch as shown. As a control for the specificity of the pulldown the histone octamer is preincubated with anti-H2A antibody (+). (D) Wild type or mutant LLD peptide was added to INO80 remodeling reactions at the indicated concentrations with or without ATP added and incubated for 30 min. 50nM of nucleosomes were incubated with 75nM of INO80 complex with addition of 80 μM ATP. The remodeling reactions were stopped, INO80 stripped off and analyzed by EMSA as shown. Three replicates are performed for each experiment and error bars represent the mean ± SD.

We next want to determine whether mutation of LLD alters the conformation of the grappler domain to indirectly interfere with Arp5 binding to the acidic pocket or the LLD region directly binds the acidic pocket and is dependent on these three residues. One of the most characterized arginine anchors is the LANA (Latency Associated Nuclear Antigen) peptide that by pulldown assays and structural studies has been shown alone to directly bind histones(51,52). We find a biotinylated 30 amino acid peptide containing the LLD region of Arp5 can bind histones slightly more efficiently than biotinylated LANA peptide and these interactions are both blocked by an antibody directed again histone H2A (Figure 6C). A biotinylated peptide containing the arginine anchor of Arp5 (R3) also binds to histones as shown by pulldown assays and is blocked by anti-H2A antibodies, comparable to that observed for the LANA peptide (Figure 6C). These data suggest that within the grappler domain of Arp5 there are two regions that can independently bind to the acidic pocket of nucleosomes.

The specificity of the interactions of the LLD peptide with nucleosomes is tested by finding if the LLD peptide at particular concentrations can interfere with nucleosomes mobilization by INO80. LLD peptide at about 28- and 84-fold higher molar ratio to nucleosomes reduced INO80 mobilization of nucleosomes by 50% and 16%, respectively (Figure 6D, lanes 5-8 and Figure S8C). A mutant version of the LLD peptide in which the LLD residues are mutated to alanine did not interfere with INO80 remodeling even at the highest concentration where wild type LLD efficiently blocks nucleosome mobilization (Figure 6D, compare lane 9-16 to 1-8, and Figure S8C). These results show Leu 567, Leu 568 and Glu 571 of Arp5 likely bind nucleosomes which can competitively block INO80 from remodeling.

### Distinct roles for the LLD and DBD regions of Arp5 in mobilizing nucleosomes by INO80

We use the same approach mentioned earlier to map DNA movement and displacement to determine at which stage(s) mutation of LLD impacts INO80 remodeling and compare that to mutation of DBD and the arginine anchor of Arp5. On the exit side INO80 initially moves DNA 20 bp but then rapidly transitions to 32 bp from its starting position (Figure S9). The LLD mutant complex moves DNA 32 bp from its starting position 3-fold slower than wild type, which is more than 2 times faster than mutant DBD or arginine anchor (Figures 7A-B and S9and Table 1). DBD and R3 mutant complexes also move approximately half as much nucleosomes as does the LLD mutant and together reflect a 4-fold difference between LLD and DBD or R3 mutants (Table 1). On the exit side, the R3 mutant is ∼10-fold more defective than DBD and LLD mutant complexes for moving DNA 20 bp from its starting position (Figure 7C-D and Table 1). The LLD and DBD mutant complexes are ∼5-fold slower for moving DNA 20 bp on the entry side of nucleosomes than wild type (Table 1). The one feature of LLD most unique to this complex is the defect in displacing DNA from the histone octamer on the entry side as seen by net loss of DNA crosslinking at H2B 53 (Figure 7E-F and S10). DNA is comparably displaced at the entry side by wild type and DBD mutant, but displacement is reduced 3 or 3.5-fold compared to the respectively the DBD mutant or wild type complex (Table 1). The R3 mutant initially displaces DNA at the entry side as efficiently as wild type, but after a short time stops presumable because of its strong defect for moving DNA at the entry and exits sides (Figures 7E-F and S10). Previously we had shown that INO80 on the entry side of nucleosomes can selectively replace H2A.Z for H2A, which is presumed to be link to INO80’s ability to displace DNA from the histone octamer and may be connected to the impact of the LLD mutation on the in vivo activity of Arp5 (13).

**Figure 7.**
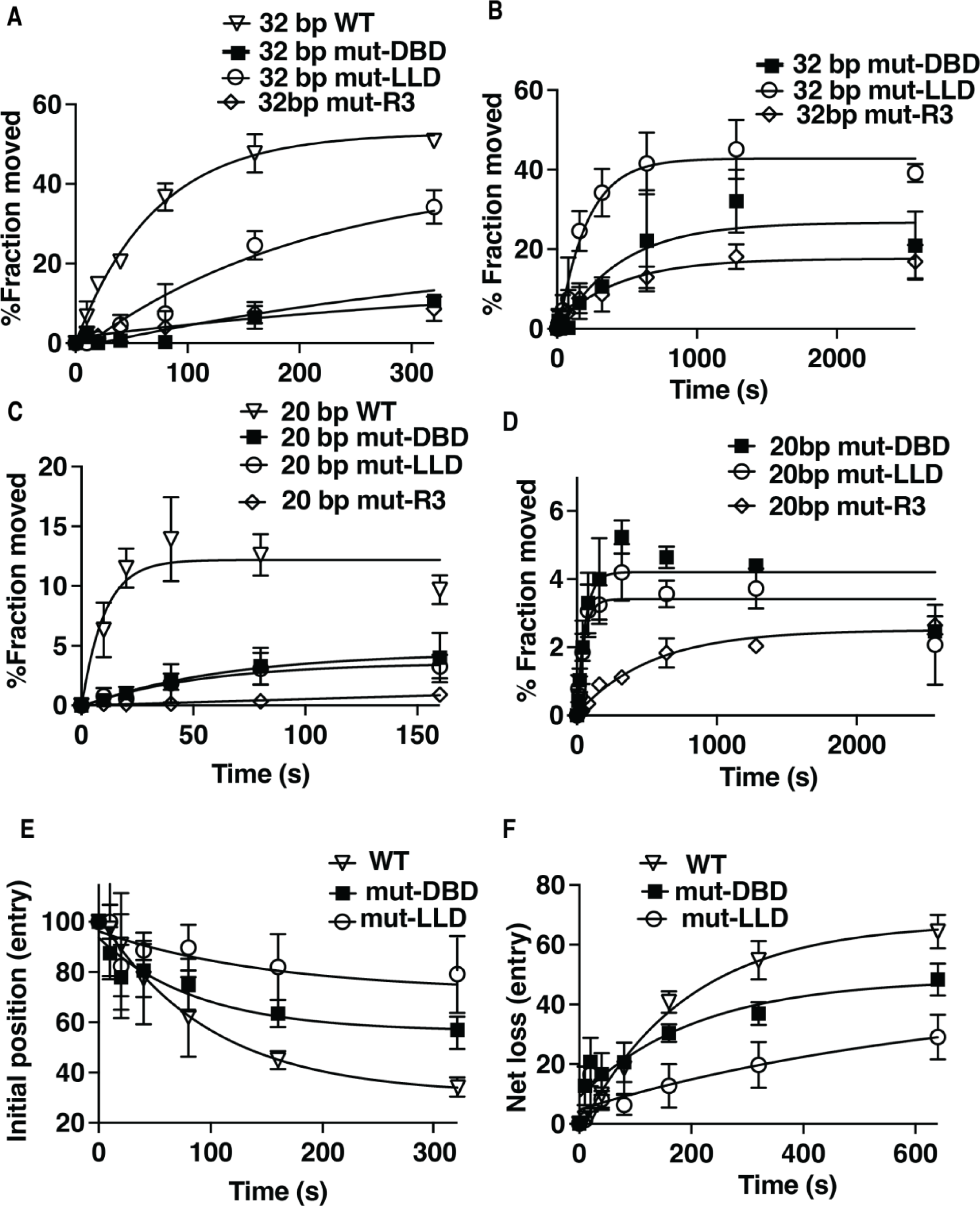
DNA displacement on the entry side is reduced when the LLD patch of Arp5 is mutated. (A-D) The extent of DNA moves (A-B) 32 bp on the exit and (D-C) 20 bp on the entry side is plotted versus time using a photoreactive probe attached to residue 53 of histone H2B for wild type (open triangle), mutant (mut)-DBD (closed square) and mutant (mut)-LLD (open circle) Arp5 complexes. The same symbols are also used in (E-F). In (C-D) the reactions are monitored for longer times. (E) The rate at which the initial DNA position disappears is shown for wild type and mutant Arp5 as in (A-D). (F) The net loss of DNA crosslinking on the entry side is plotted versus time similar to (A-D). Three replicates are performed for each condition and error bars represent the mean ± SD.

## Discussion

In this study, we find evidence for INO80 from *Saccharomyces cerevisiae* engaging nucleosomes in two distinct ways that have differing outcomes in vivo involving the grappler domain of the Arp5 subunit. Two contiguous regions of Arp5’s grappler domain are found to be associated with the acidic pocket of nucleosomes by crosslinking Arp5 to a site near the top of the pocket. A cluster of three arginines suggested by structural modeling of Arp5 to be an arginine anchor that binds the acidic pocket of nucleosomes are adjacent to one of these two regions(33,34). Mutation of the three arginines to alanine reduces the rate at which INO80 mobilizes nucleosomes like that shown previously for *Chaetomium thermophilium* INO80(34). We provide direct evidence for these arginines being required for Arp5 binding the acidic pocket by site-specific histone crosslinking. Surprisingly, the arginine anchor mutant version of Arp5 in vivo does not display the same phenotype as when Arp5 is absent under DNA replication or transcription elongation stress or for repairing DNA damage caused by the alkylating agent methyl methansulfonate. The in vivo data suggest there may be an activity other than mobilizing nucleosomes that requires INO80 and its Arp5 subunit in these circumstances.

We find by scanning for conserved amino acids in both regions crosslinked to the acidic pocket that only mutations at the junction of the two regions show sensitivity to DNA damage and DNA replication and transcription elongation stress similar to that when Arp5 is absent. Mutating leucine and glutamate in this region to alanine blocks binding of Arp5 at the acidic pocket, while not disrupting Arp5 binding at an adjacent region on the histone octamer surface close to SHL2. We find that a short peptide of this region alone is capable of binding to histone octamers much like arginine anchors from Arp5 or the prototypical one from the LANA protein. This same peptide also interferes with INO80 mobilizing nucleosomes by presumably competing for Arp5 binding to the acidic pocket and is eliminated when the two leucines and one glutamate are mutated to alanine. The hydrophobic nature of this region resembles the hydrophobic patch of Swc6 that binds to the acidic pocket of nucleosomes in the SWR1 complex, which promotes dimer exchange (53,54). The region of H2A that is contacted by this hydrophobic patch of Swc6 is where differences between H2A and H2A.Z (Htz1) may contribute to the specificity of SWR1 to replace H2A with H2A.Z(54).

Based on the cross configuration of the grappler domain, the LLD region is far removed from the arginine anchor and acidic pocket of nucleosomes. This difference in locations suggest the arginine anchor and LLD patch likely bind mutually exclusive of each other at the acidic pocket. The region of Arp5 crosslinked to residue 80 of histone H3 provides additional evidence for a grappler domain conformation different from the cross configuration, because it is incongruent with the cryo-EM/AlphaFold2 cross grappler structure(8). The Arp5 interactions detected near residue 80 of H3 are dependent on the LLD region binding to the acidic pocket, seen only by histone crosslinking when both the LDD and DBD are mutated. These data suggest the interactions of Arp5 detected by histone crosslinking at H3 residue 80 are likely part of the same Arp5 conformation where the LLD binds the acid pocket. The other Arp5 grappler structure observed by cryo-EM referred to as the parallel conformation is also not congruent with the LLD binding to the acidic patch because although the brace domain of Arp5 shifts dramatically towards the dyad axis, the arginine anchor remains bound at the acidic pocket the same as the cross configuration(29). These data together suggest binding of LLD to the acidic pocket and the N-terminal part of the grappler near residue 80 of H3 are part of a conformation of Arp5 not previously identified.

Distinct modes of INO80 chromatin remodeling activity are coupled to the different ways in which the grappler domain can bind to the acidic pocket of nucleosomes. Besides positioning nucleosomes in a DNA shape-specific manner and spacing nucleosomes, INO80 also replaces H2A for H2A.Z as shown both in vitro and in vivo(13,21–23,25). The requirement of the LLD region of Arp5 for INO80 to relieve DNA replication stress implicates the LLD region is involved in the dimer exchange activity of INO80 in contrast to the arginine anchor of Arp5. H2A.Z is crucial for regulating initiation of early replication origins and replication timing (55). The H2A.Z-H2B dimer is switched for H2A-H2B on the newly synthesized DNA strand and is required for maintaining genome integrity (56). INO80 also helps resolve a form of replication stress called R-loops that occur when replication forks collide with transcription by removing H2A.Z(7,57,58). Our biochemical data shows the mutant LLD represses the nucleosome mobilizing activity of INO80 less than the mutant arginine anchor and points to the observed in vivo phenotype for the LLD mutant not likely being due to its reduced nucleosome mobilizing activity. We find by tracking DNA movement and displacement that the LLD mutant INO80 complex is most defective in displacing DNA from the entry side of nucleosomes proximal to the long linker DNA. INO80 has been shown to preferentially replace H2A.Z-H2B dimers on this side of nucleosomes for H2A-H2B by 3-color single molecule FRET assays(13). Together our data suggest the LLD mutation targets the dimer displacement activity of INO80, which may prove useful to examine in vivo the role of INO80’s dimer exchange activity separate from its nucleosome mobilizing activity.

## Author Contributions

JG and SP performed the site-directed crosslinking of histones to INO80 and SP performed the site-directed crosslinking of DNA to INO80 and peptide mapping. JG did the peptide competition experiment with INO80 remodeling and YH did the peptide pulldown with histone octamers. Purification of wild type and mutant Arp5 complexes along with binding and gel shift experiments were performed by JG, SP and SS. The Anchor Away experiments were performed by YH and KB and the plasmids used for complementation assays with wild type and mutant Arp5 constructed by YH. SP constructed and characterized the truncated forms of Arp5. SS did all the mapping of DNA movement and displacement experiments. SS did most of the figure preparation with assistance from JG and SP.

## Tables

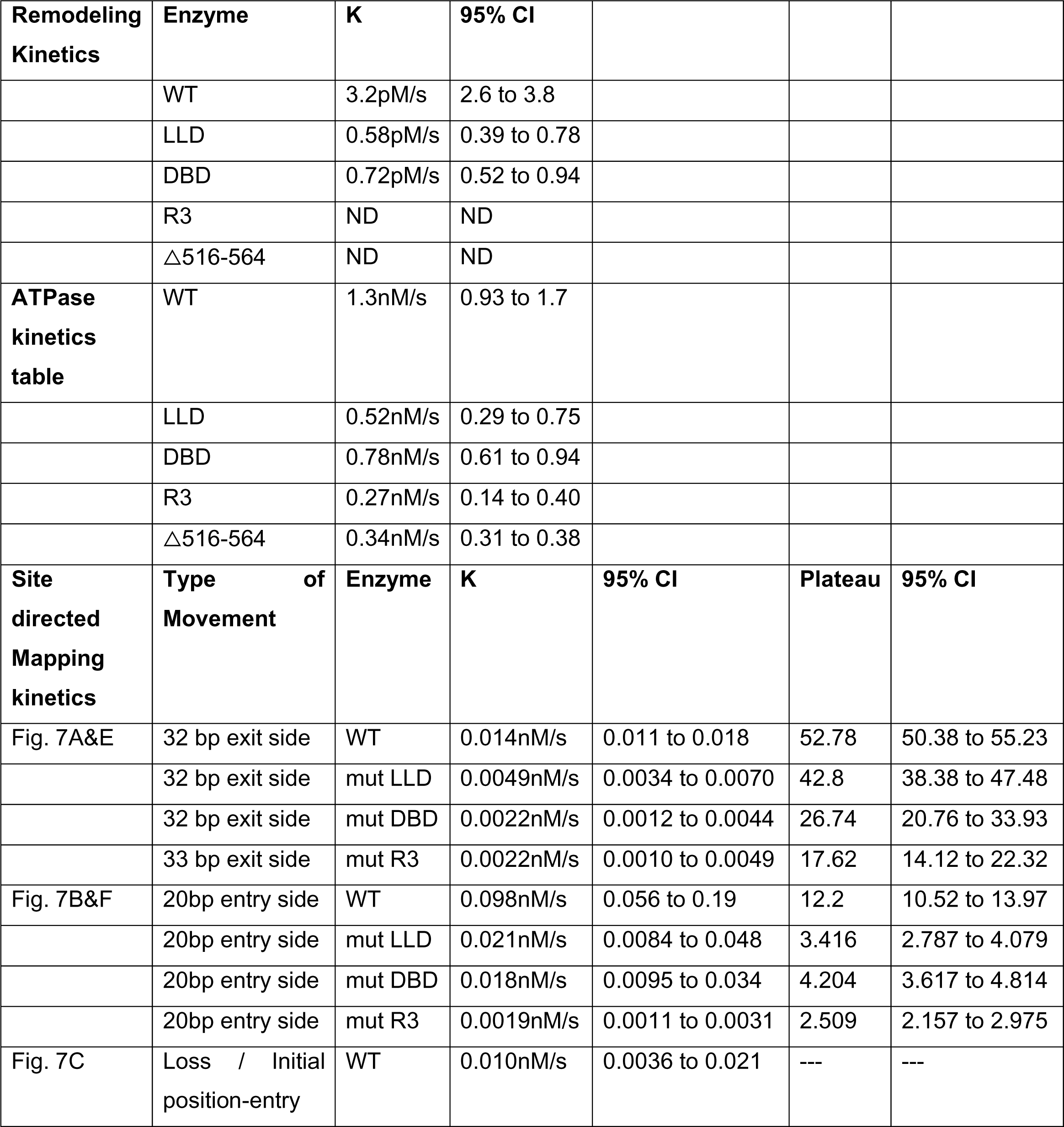
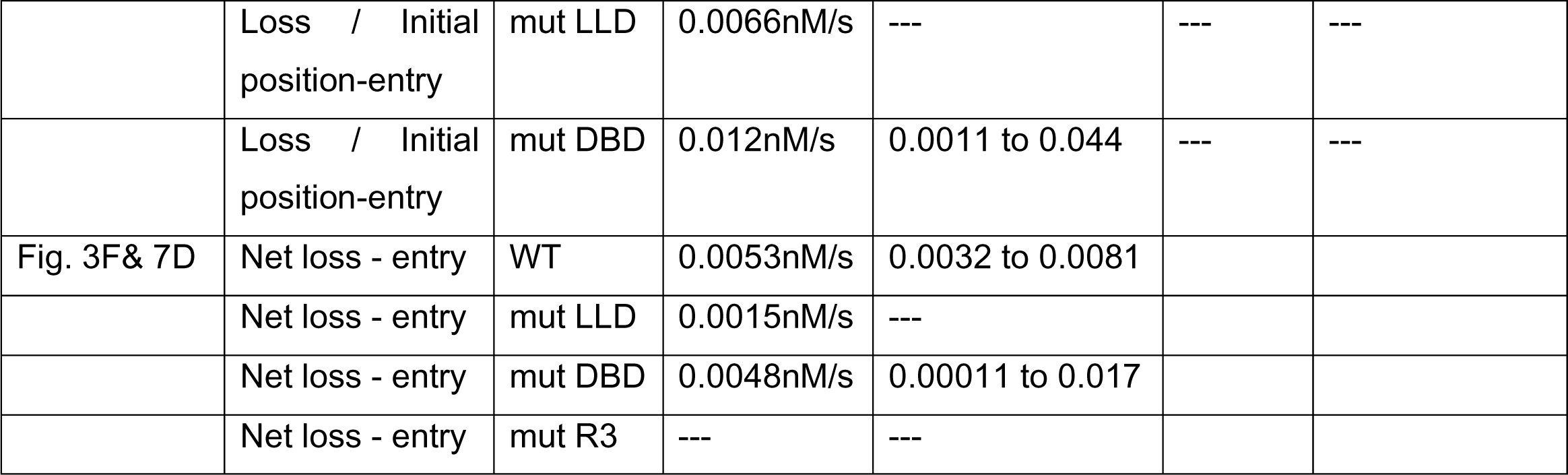

| **Remodeling Kinetics** | **Enzyme** | **K** | **95% CI** |  |  |  |
| --- | --- | --- | --- | --- | --- | --- |
|  | WT | 3.2pM/s | 2.6 to 3.8 |  |  |  |
|  | LLD | 0.58pM/s | 0.39 to 0.78 |  |  |  |
|  | DBD | 0.72pM/s | 0.52 to 0.94 |  |  |  |
|  | R3 | ND | ND |  |  |  |
|  | △516-564 | ND | ND |  |  |  |
| **ATPase kinetics table** | WT | 1.3nM/s | 0.93 to 1.7 |  |  |  |
|  | LLD | 0.52nM/s | 0.29 to 0.75 |  |  |  |
|  | DBD | 0.78nM/s | 0.61 to 0.94 |  |  |  |
|  | R3 | 0.27nM/s | 0.14 to 0.40 |  |  |  |
|  | △516-564 | 0.34nM/s | 0.31 to 0.38 |  |  |  |
| **Site directed Mapping kinetics** | **Type of Movement** | **Enzyme** | **K** | **95% CI** | **Plateau** | **95% CI** |
| Fig. 7A&E | 32 bp exit side | WT | 0.014nM/s | 0.011 to 0.018 | 52.78 | 50.38 to 55.23 |
|  | 32 bp exit side | mut LLD | 0.0049nM/s | 0.0034 to 0.0070 | 42.8 | 38.38 to 47.48 |
|  | 32 bp exit side | mut DBD | 0.0022nM/s | 0.0012 to 0.0044 | 26.74 | 20.76 to 33.93 |
|  | 33 bp exit side | mut R3 | 0.0022nM/s | 0.0010 to 0.0049 | 17.62 | 14.12 to 22.32 |
| Fig. 7B&F | 20bp entry side | WT | 0.098nM/s | 0.056 to 0.19 | 12.2 | 10.52 to 13.97 |
|  | 20bp entry side | mut LLD | 0.021nM/s | 0.0084 to 0.048 | 3.416 | 2.787 to 4.079 |
|  | 20bp entry side | mut DBD | 0.018nM/s | 0.0095 to 0.034 | 4.204 | 3.617 to 4.814 |
|  | 20bp entry side | mut R3 | 0.0019nM/s | 0.0011 to 0.0031 | 2.509 | 2.157 to 2.975 |
| Fig. 7C | Loss / Initial position-entry | WT | 0.010nM/s | 0.0036 to 0.021 | --- | --- |
|  | Loss / Initial<br>position-entry | mut LLD | 0.0066nM/s | --- | --- | --- |
|  | Loss / Initial<br>position-entry | mut DBD | 0.012nM/s | 0.0011 to 0.044 | --- | --- |
| Fig. 3F& 7D | Net loss - entry | WT | 0.0053nM/s | 0.0032 to 0.0081 |  |  |
|  | Net loss - entry | mut LLD | 0.0015nM/s | --- |  |  |
|  | Net loss - entry | mut DBD | 0.0048nM/s | 0.00011 to 0.017 |  |  |
|  | Net loss - entry | mut R3 | --- | --- |  |  |

