## Supplementary File for "Conformational switching of Arp5 subunit differentially regulates INO80 chromatin remodeling"

**Supplementary Figures S1-S10, Table S1.**

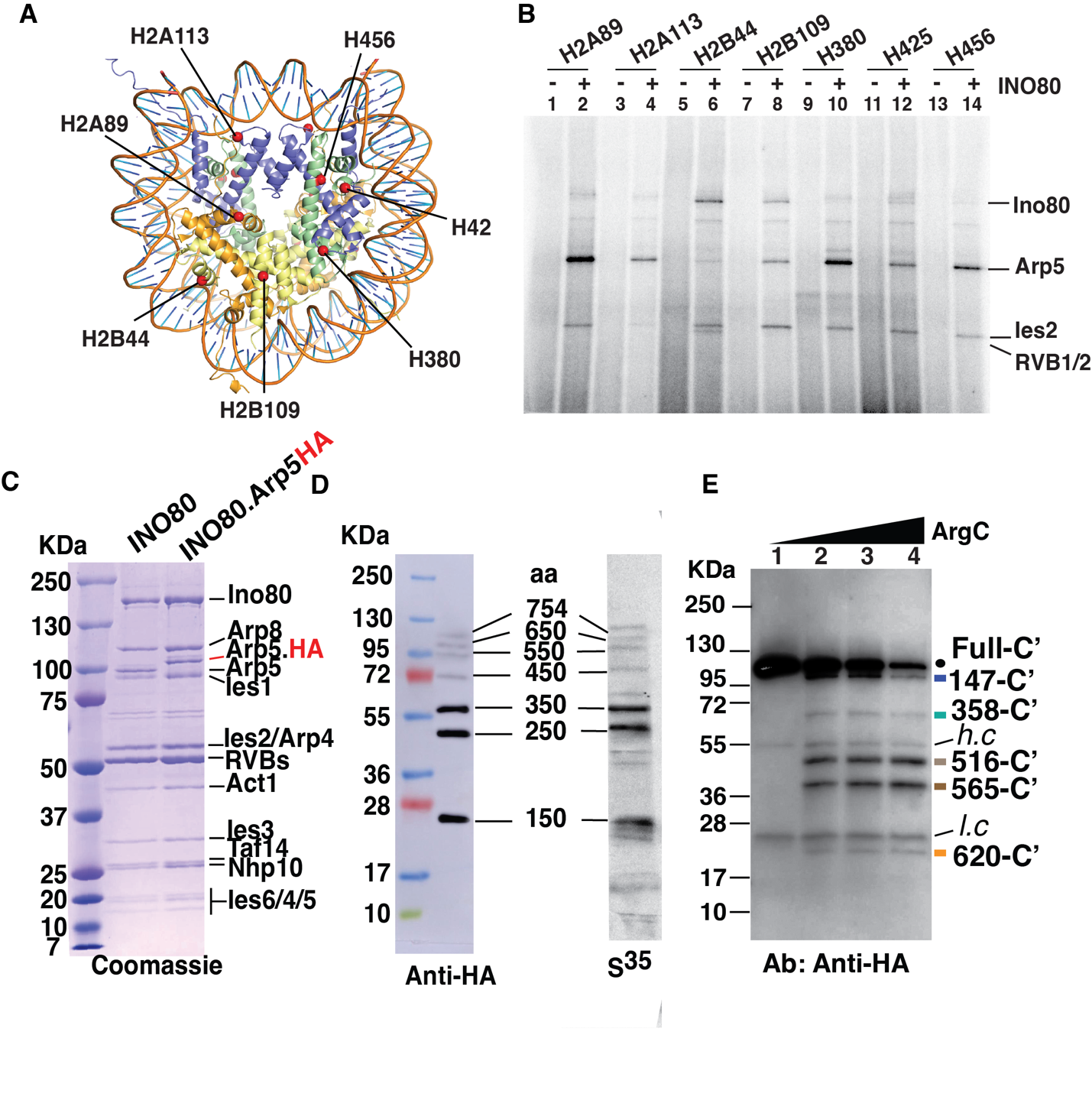

**Figure S1. Parts of Arp5 associated with the acid pocket of nucleosomes are required for the remodeling activity of INO80.**

(A) The positions in the nucleosome scanned by site-specific histone crosslinking are shown. (B) The phosphorimage shown is representative of the data graphed in Figure 1B. (C) Purified wild type INO80 and the Arp5 HA tagged version of INO80 were analyzed on a 4-20% SDS-PAGE and stained with Coomassie blue. (D) The S^35^ labeled Arp5 protein molecular weights used to determine the fragment sizes of proteolytically cleaved photocrosslinked Arp5 are shown. (E) Labeled proteolytic fragments of Arp5 are separated on a 4-20% SDS-PAGE and visualized by Western Blotting anti-HA antibody. As shown in Fig.1C.

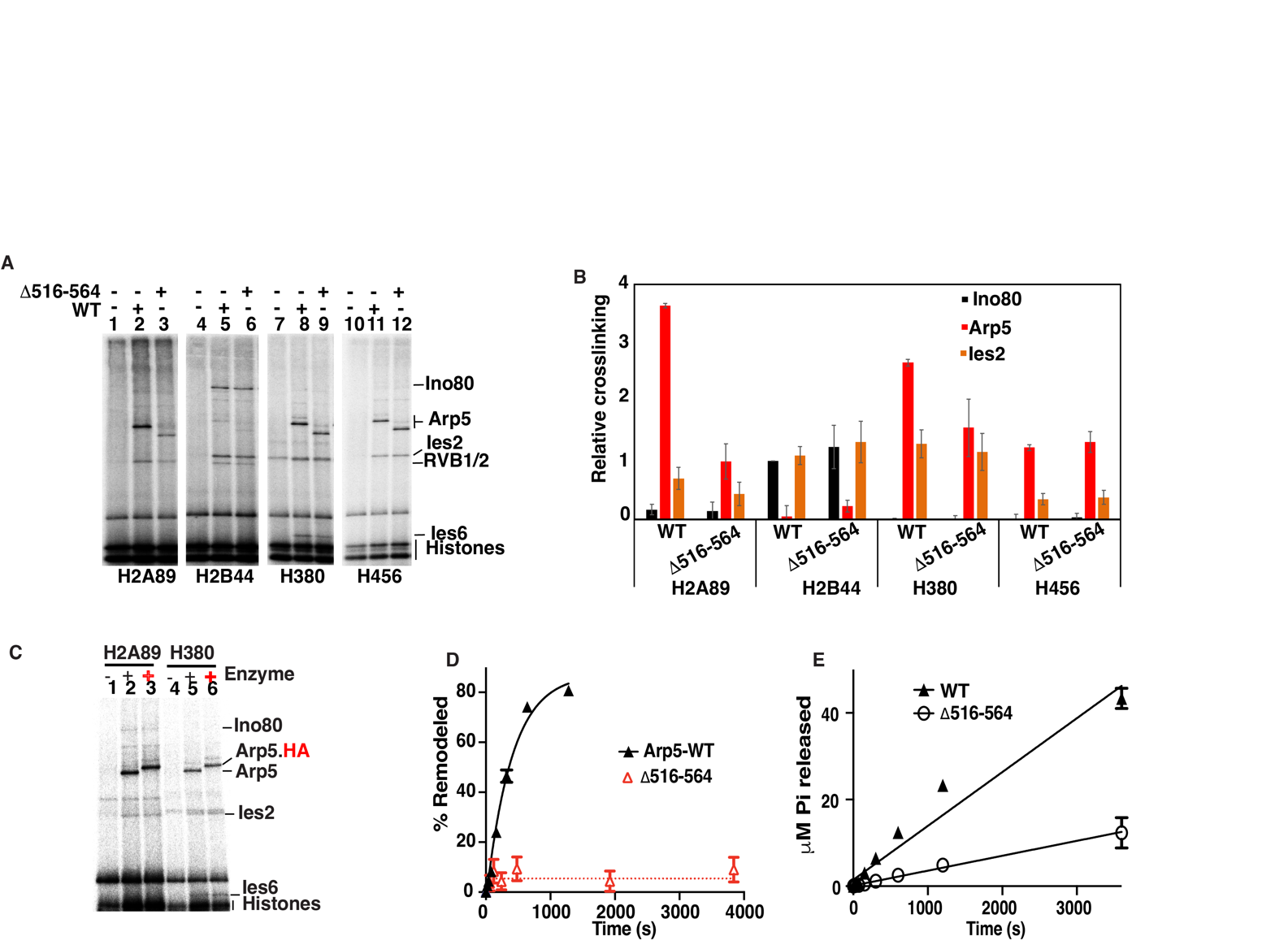

**Figure S2.** (A-B) The interactions of Arp5 with the histone octamer face of nucleosomes are probed by site-specific histone crosslinking for WT and Arp5Δ516-564. The relative efficiency of crosslinking Arp5, Ies2 and Ies6 is shown for three replicates with Three replicates are performed for each experiment and error bars represent the mean ± SD. (C) Phosphorimage of SDS-PAGE showing the crosslinking of WT Arp5 and WT Arp5.HA at H2A89 and H380 octamer positions. (D) The remodeling activity of the Arp5Δ516-564 INO80 complex is shown from 3 replicates± SD like that shown in Figure 1F. (E) The ATPase activity of mutant and wild type INO80 was assayed under the same conditions as in (D).

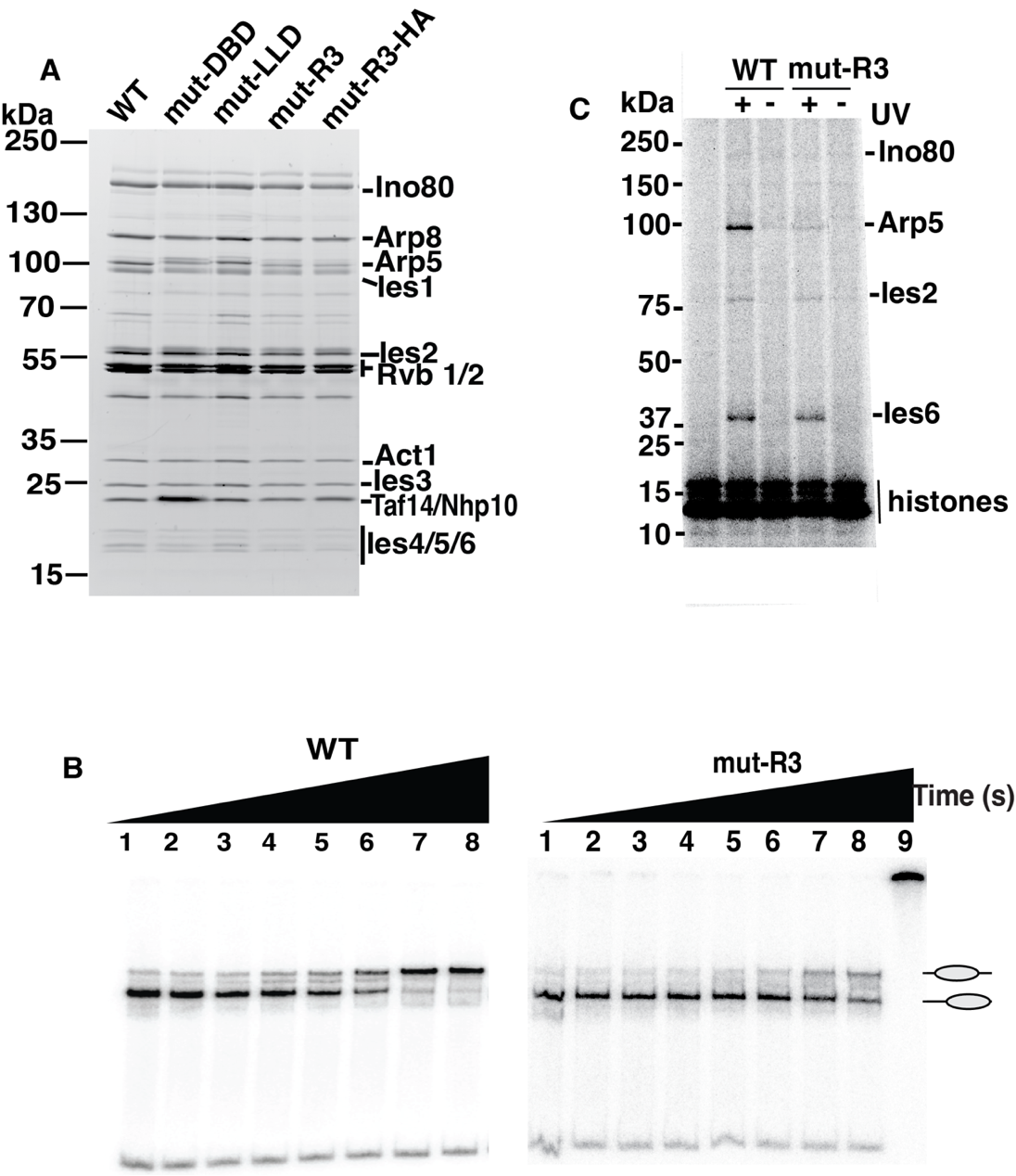

**Figure. S3**

(A) SDS-PAGE 4-20% profile of the WT, LLD, DBD, R3, R3+HA tag mutant complexes of INO80. (B)The nucleosome remodeling activity of wild type (WT) and mutant (R3) INO80 was measured by EMSA. Nucleosomes (50nM) were incubated with of INO80 complex (75nM) with addition of 80 μM ATP (C) Histone crosslinking is performed for wild type and R3 mutant Arp5 R3 WT entry exit containing INO80 with samples being either irradiated (+) or not (-).

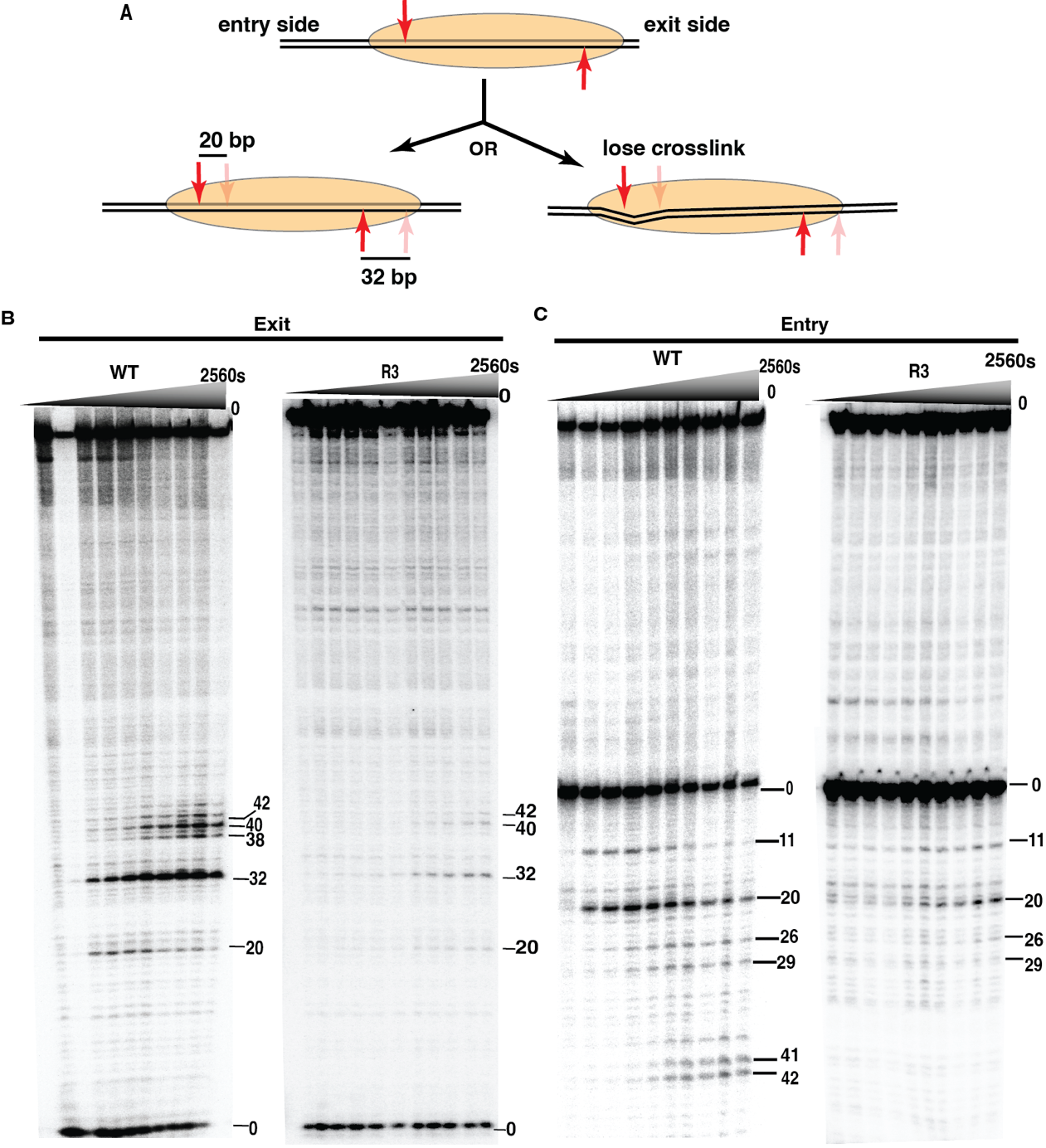

**Figure. S4** (A) The approach for mapping changes in histone-DNA contacts at the entry and exit side of nucleosome is shown for side-directed crosslinking and how it can measure DNA movement and displacement. (B-C) A phosphorimage of the 6% denaturing polyacrylamide used to examine DNA movement and translocation on the entry and exit side is shown that is representative of that used in the graphs shown in Figure 3D-F. For each experiment WT or mutants, 80nM of nucleosomes were incubated with 80nM of INO80 complex with addition of 80 μM ATP.

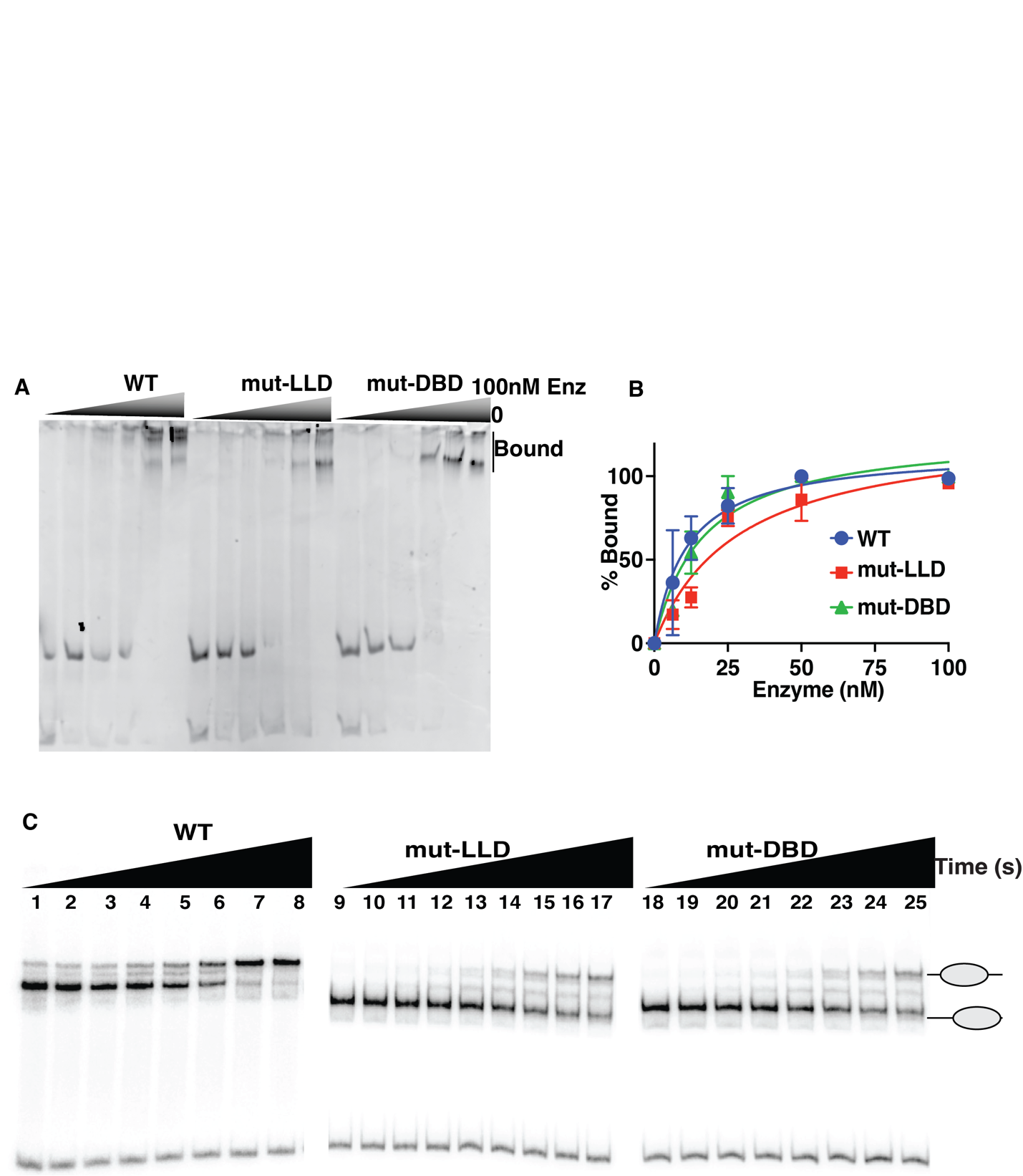

**Figure.S5**

(A) INO80 binding assays for WT, LLD and DBD with 601 nucleosomes (25 nM) are determined by EMSA with 0-100 nM INO80. (B). The binding data in (A) is plotted with INO80 concentration versus the percent of nucleosomes bound error bars represent ± SD (C). The nucleosome remodeling activity of wild type (WT) and mutant (LLD or DBD) INO80 was measured by EMSA. 50nM of nucleosomes were incubated with 75nM of INO80 complex with addition of 80 μM ATP

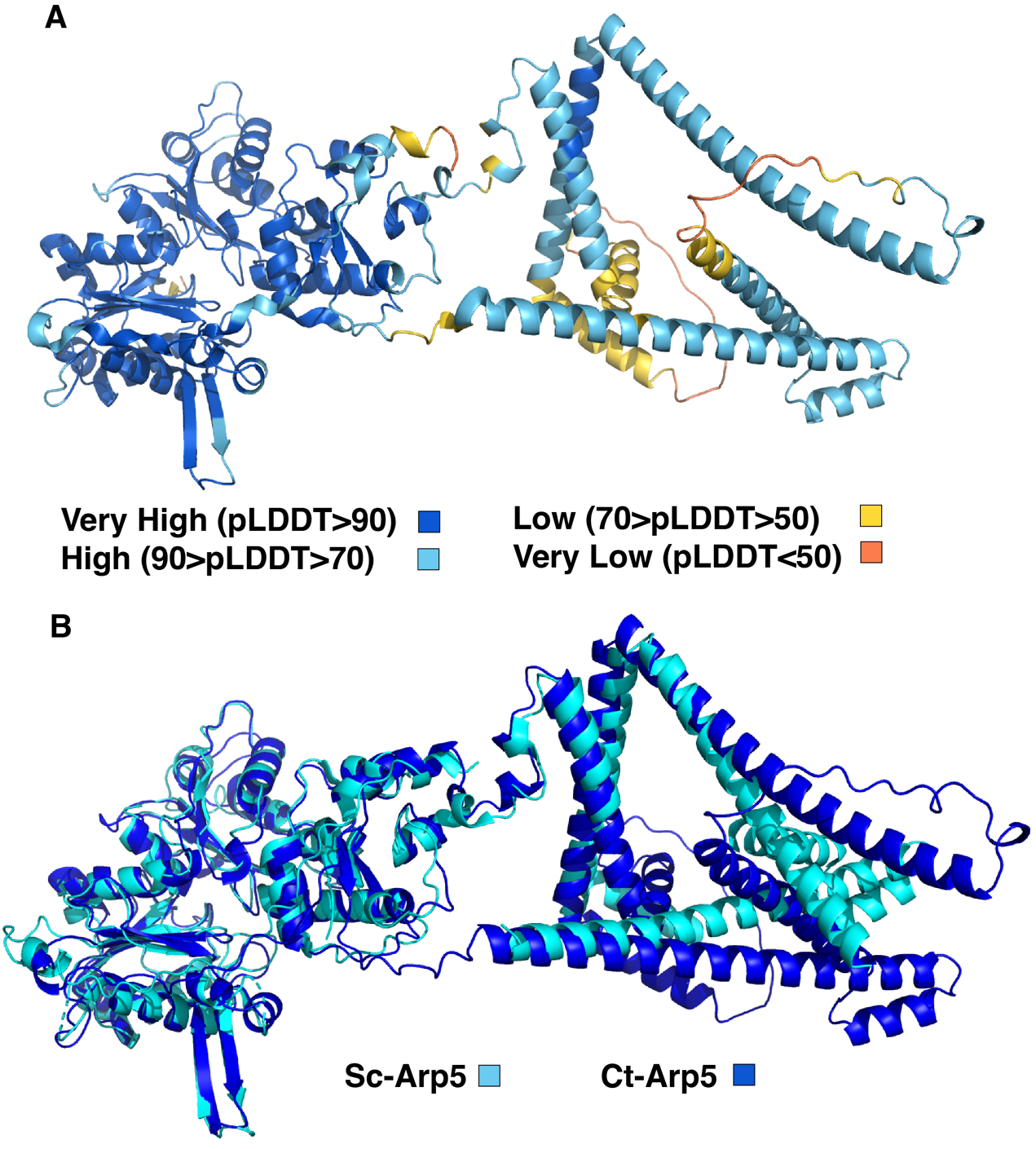

**Figure S6.**

**(A)** Model of Arp5 generated using AlphaFold2, colors highlight the confidence of residue structures calculated as pLDDT (predicted local distance difference test). (B) Model overlay of *S. cerevisiae* and *C.thermophila* Arp5 generated using AlphaFold2.

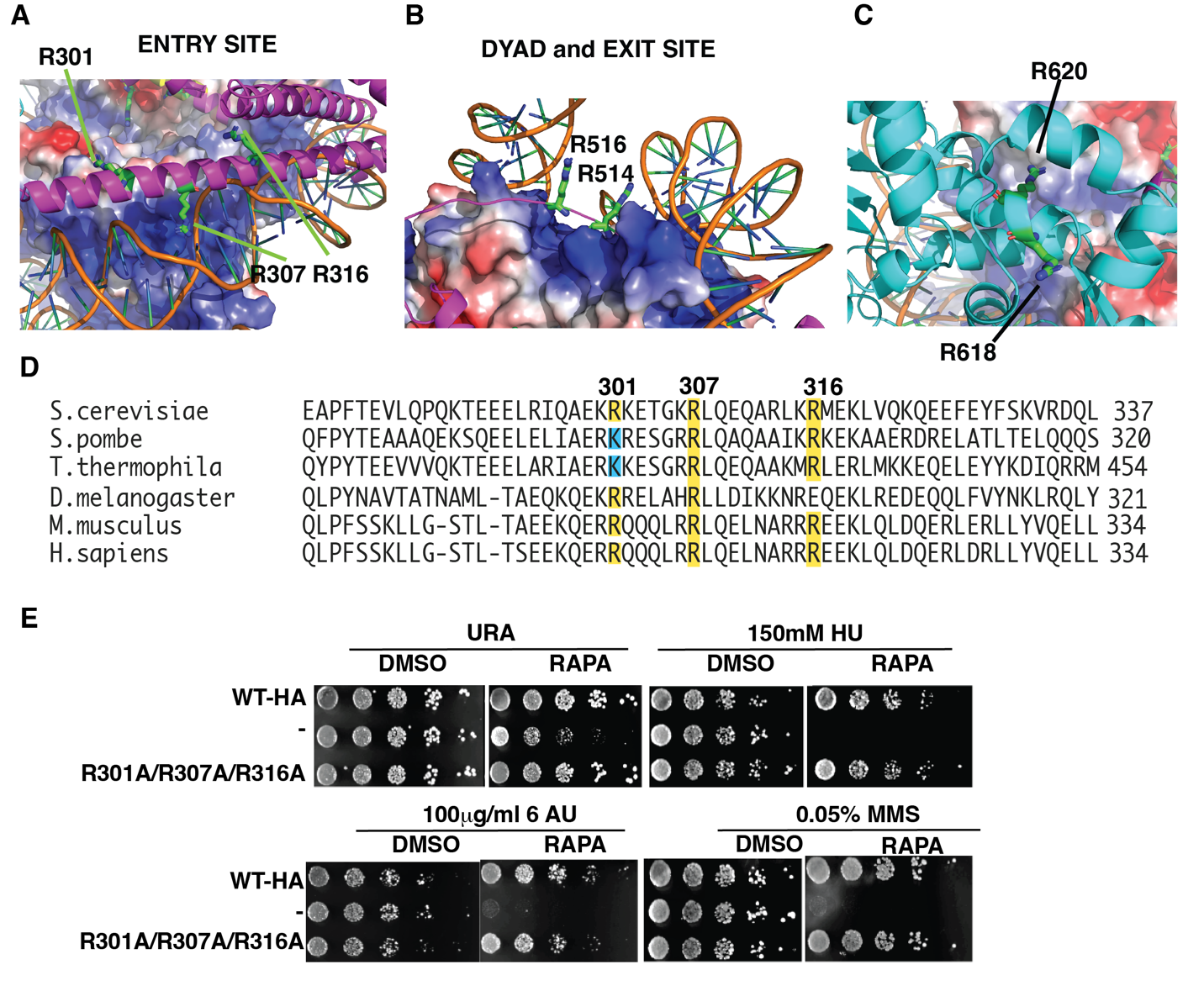

**Figure S7.**

Structural model of Arp5 shows the location of (A-D) various arginine’s that were tested in spot assays to determine their impact on the in vivo activity of Arp5 as shown in (E).

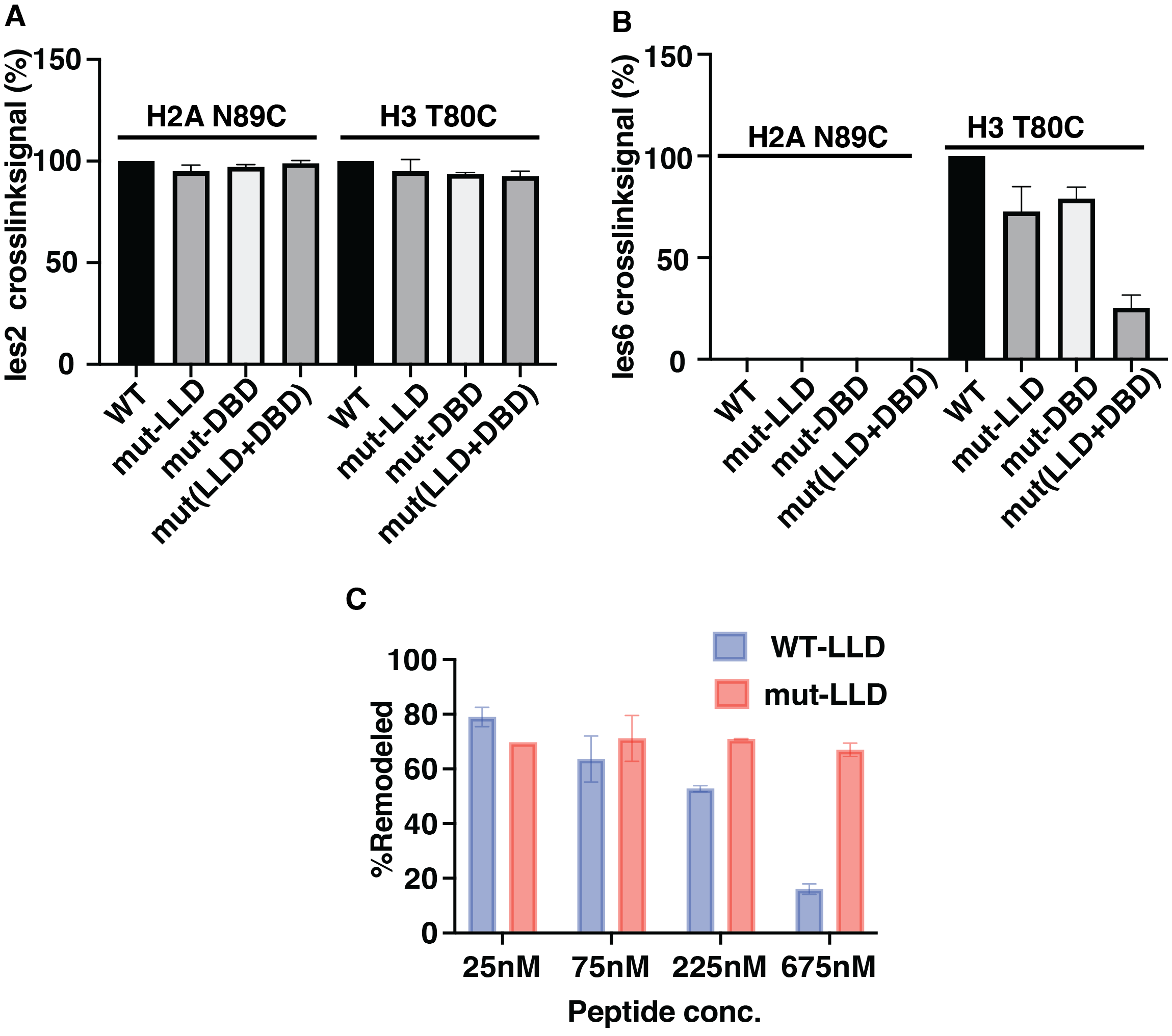
**Figure.S8**

(A-B) Binding of Ies2 and Ies6 near the acid pocket (H2A N89C) and another lateral position on the histone octamer (H3 T80C) is probed by site-specific histone crosslinking for wild type (WT) and mutant INO80 (LLD and DBD). (C) Peptide competition data in Figure 4D plotted for the peptides used in the assay wild type (LLD) and mutant peptide. Data shown is from 3 replicates± SD.

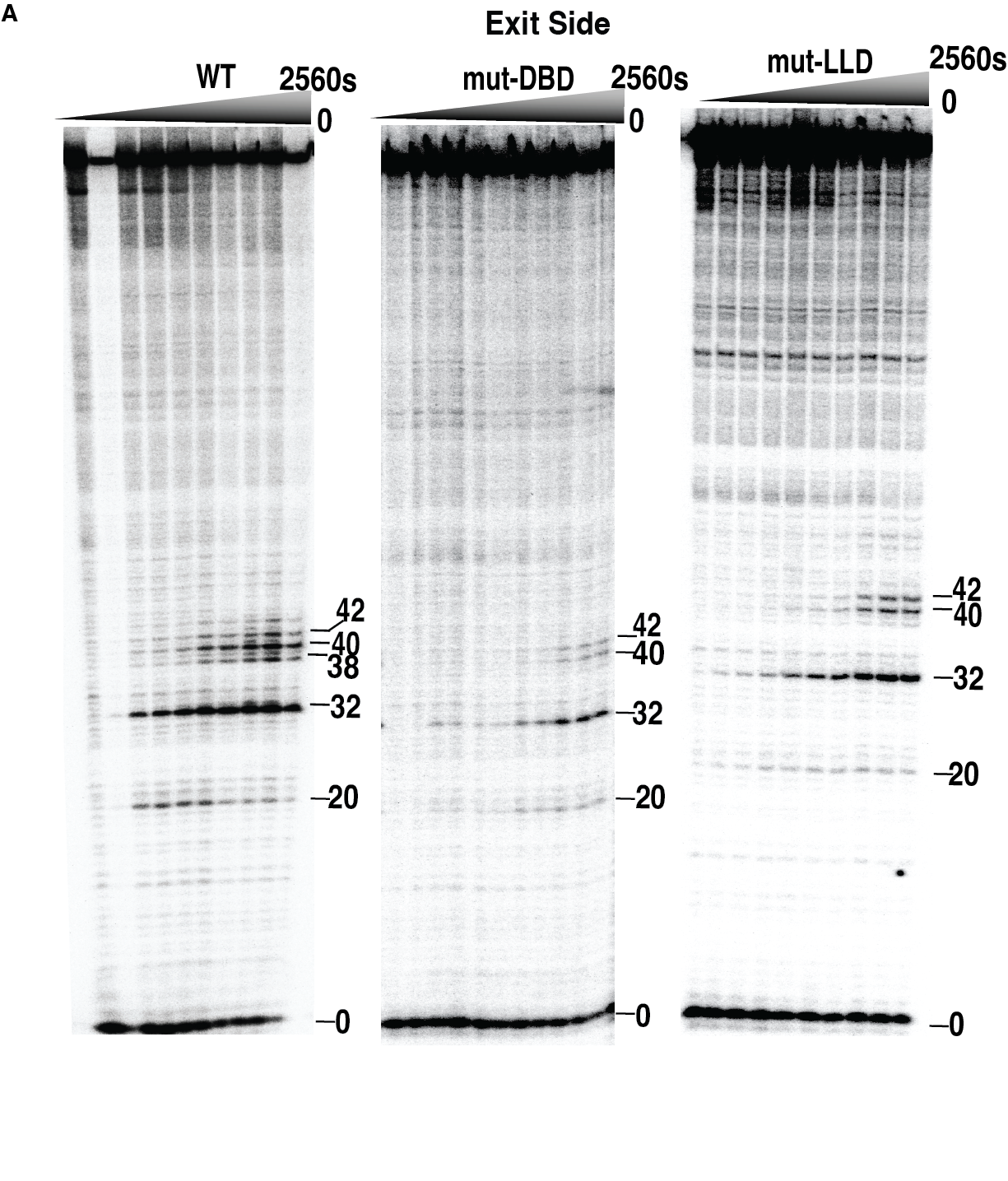

**Figure S9. Site-directed mapping on the exit side for wild type and DBD and LLD mutant Arp5 containing INO80 complexes**

A phosphorimage of the 6% denaturing polyacrylamide used to examine DNA movement and translocation on the exit side is shown that is representative of that used in the graphs shown in Figure 7A-F. Nucleosomes (80nM) were incubated with INO80 complex (80nM) with addition of 80 μM ATP

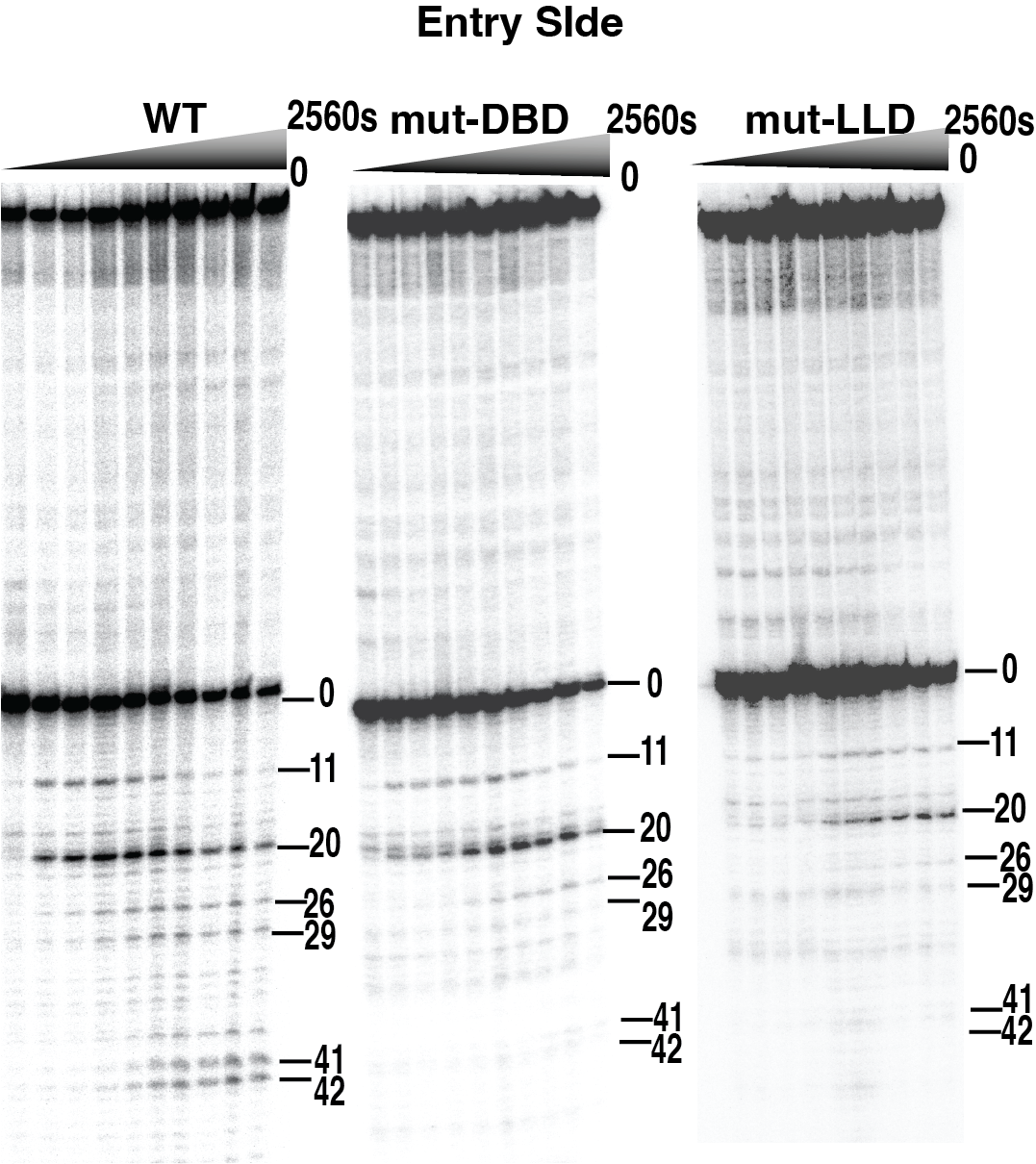

**Figure S10. Site-directed mapping on the entry side for wild type and DBD and LLD mutant Arp5 containing INO80 complexes**

A phosphorimage of the 6% denaturing polyacrylamide used to examine DNA movement and translocation on the entry side is shown that is representative of that used in the graphs shown in Figure 7A-F. Nucleosomes (80nM) were incubated with INO80 complex (80nM) with addition of 80 μM ATP

**Table S1**

| **REAGENT or RESOURCE** | **SOURCE** | **IDENTIFIER** |
| --- | --- | --- |
| **Antibodies** | | |
| Anti-HA | Invitrogen | cat# 26183 |
| Anti-H2A | Millipore | cat#07-146 |
| **Yeast strains** | | |
| INO80-FLAG, ((BY4741) MATα his3Δ1 leu2Δ0 met15Δ0) | Brahma et. al, 2018 (1) | N/A |
| Arp5HA, ((BY4741) MATα his3Δ1 met15Δ0 Ino80::Ino80-2XFLAG Arp5Δ::ARP5-HA) | This study | N/A |
| Arp8ΔN, isogenic to INO80-FLAG except Arp8::Arp8ΔN | Brahma et. al, 2018 (1) | N/A |
| WT-W303, ((CY1827) MATα tor1-1 fpr1::loxP-LEU2-loxP RPL13A 2×FKBP12::loxP bar1∆::HISG) | Gift from Craig Peterson (2) | N/A |
| Arp5-FRB-W303, isogenic to WT-W303 except ARP5-FRB::KanMX | This study | N/A |
| ΔArp5-XS, ((BY4741) MATα his3Δ1 leu2Δ0 met15Δ0 ura3Δ0 Arp5Δ) | Gift from Xuetong Shen | N/A |
| ΔArp5, isogenic to ΔArp5-XS INO80-2XFLAG::Leu2 | This study | N/A |
| Arp5-WT, isogenic to ΔArp5 except Arp5::URA3 | This study | N/A |
| B/Arp5Δ516-564, isogenic to ΔArp5 except Arp5Δ516-564::URA3 | This study | N/A |
| B/Arp5Δ565-620, isogenic to ΔArp5 except Arp5Δ565-620::URA3 | This study | N/A |
| B/Arp5R3, isogenic to ΔArp5 except Arp5R482A, R488A,R496A::URA3 | This study | N/A |
| B/Arp5LLD, isogenic to ΔArp5 except Arp5 L567A,L568A, D571A::URA3 | This study | N/A |
| B/Arp5DBD, isogenic to ΔArp5 except Arp5 K69A, R71A, R73A, K77A, R93A, R97A::URA3 | This study | N/A |
| B/Arp5LLD+DBD, isogenic to ΔArp5 except Arp5 K69A, R71A, R73A, K77A, R93A, R97A, L567A,L568A, D571A::URA3 | This study | N/A |
| W/Arp5Δ516-564, isogenic to Arp5-FRB-W303 except Arp5Δ516-564::URA3 | This study | N/A |
| W/Arp5Δ565-620, isogenic to Arp5- FRB-W303 except Arp5Δ565-620::URA3 | This study | N/A |
| W/Arp5R3, isogenic to Arp5- FRB-W303 except Arp5 R482A, R488A,R496A::URA3 | This study | N/A |
| W/Arp5LLD, isogenic to Arp5- FRB- W303 except Arp5 L567A,L568A, D571A::URA3 | This study | N/A |
| W/Arp5DBD, isogenic to Arp5- FRB- W303 except Arp5 K69A, R71A, R73A, K77A, R93A, R97A::URA3 | This study | N/A |
| W/Arp5LLD+DBD, isogenic to Arp5- FRB- W303 except Arp5 K69A,R71A,R73A,K77A,R93A,R97A, L567A,L568A,D571A::URA3 | This study | N/A |
| W/Arp5R514,516, isogenic to Arp5- FRB- W303 except Arp5 R514A,R516A::URA3 | This study | N/A |
| W/Arp5R565, isogenic to Arp5- FRB- W303 except Arp5 R565A::URA3 | This study | N/A |
| W/Arp5R596,599, isogenic to Arp5- FRB- W303 except Arp5 R596A,R599A::URA3 | This study | N/A |
| W/Arp5R618,620, isogenic to Arp5- FRB- W303 except Arp5 R618A,R620A::URA3 | This study | N/A |
| W/Arp5 FWY, isogenic to Arp5- FRB- W303 except Arp5 F531A,W538A,Y541A::URA3 | This study | N/A |
| W/Arp5 DDD, isogenic to Arp5- FRB- W303 except Arp5 D529A,D535A,D537A::URA3 | This study | N/A |
| W/Arp5 EEE, isogenic to Arp5- FRB- W303 except Arp5 E550A,E551A,E554A::URA3 | This study | N/A |
| **Chemicals, peptides, and recombinant proteins** | | |
| Yeast Extract | Fisher Scientific | cat#BP14222 |
| Peptone | Research Products International, Corp | cat#P20250 |
| Dextrose | Fisher Scientific | cat#D16-10 |
| Adenine sulfate | Gibco | cat#25030-081 |
| Yeast Nitrogen Base | Gibco | cat#11360-070 |
| Anti-FlagM2-Beads | Gibco | cat#11140-050 |
| 2-mercaptoethanol | Sigma | cat#M6250 |
| Hydroxyurea | Sigma-Aldrich | cat# H8627 |
| Azauracil | Stem cell technology | cat#72184 |
| MMS (methyl methanesulfonate) | Stem cell technology | cat#72054 |
| Arginine C protease | Promega | cat#V1881 |
| BSA | Sigma | cat#A1470 |
| Amicon Ultra Filters | EMD Millipore | cat#UFC803024 |
| ATP,{γ-32P}-3000Ci/mmol 10mCi/ml EasyTide Lead,500 uCi | PerkinElmer Inc | cat#NEG502A500UC |
| Iodine-125 Radionuclide, 10mCi,Specific Activity: 17Ci/mg,0.1M NaOH(ph12-14), Concentration >350mCi/ml | PerkinElmer Inc | cat#NEZ033H010MC |
| Polyethyleneimine Cellulose plates | J.T Baker | cat#02-002-562 |
| PEAS | Thermofisher | cat#P6317 |
| Dynabeads MyOne Streptavidin T1 | Invitrogen | cat#65601 |
| PBS | Sigma | cat# P3813 |
| Gelcode Blue stain | Thermofisher | cat#24590 |
| ATP Solution, Tris buffered | Thermofisher | cat#R1441 |
| Pyrrolidine | Sigma | cat#394238 |
| AntiFoam agent | Sigma | cat# A8311 |
| Rapamycin | Calbiochem | cat#553210 |
| DMSO | Sigma | cat#D8418 |
| Sypro Ruby Stain | Invitrogen | cat#S12000 |
| Bacto Agar Soldifying Agent | BD Diagnostics- | cat# 214030, |
| ANTI-FLAG(R) M2 Affinity Gel | Sigma-Aldrich | cat# A2220 |
| IPTG | Sigma | cat#16758 |
| SuperSignal™ West Femto Maximum Sensitivity Substrate | Thermofisher | cat#34094 |
| Anti-HA Agarose beads, 50% slurry | Pierce | cat#26182 |
| Critical commercial assays | | |
| TNT Quick Coupled Transcription/Translation System | Promega | cat#31980-030 |
| Plasmids | | |
| pFA6a-FRB-KanMX6 | EUROSCARF | cat#P30578 |
| pRS-416 | Sikorski RS et. al,(2) | N/A |
| pRS416 Arp5 | Brahma et.al,(1) | N/A |
| **Peptides** | | |
| WT LLD: Biotin-PEAFEEALEYEYKDIVELERLLLEHDPNFT) | CPC scientific Inc | N/A |
| Mutant LLD: Biotin-PEAFEEALEYEYKDIVELERLAAEHAPNFT) | CPC scientific Inc | N/A |
| WT LANA (Biotin-MAPPGMRLRSGRSTGAPLTRGSC) | CPC scientific Inc | N/A |
| Mutant LANA (Biotin-MAPPGMALASGASTGAPLTRGSC) | CPC scientific Inc | N/A |
| **Recombinant DNA** | | |
| Arp5DBD | This study | N/A |
| Arp5LLD | This study | N/A |
| Arp5EEE-HA | This study | N/A |
| Arp5FWY | This study | N/A |
| Arp5FWY+DDD | This study | N/A |
| Arp5R565A | This study | N/A |
| Arp5R596A/R599A | This study | N/A |
| Arp5R618A/R620A | This study | N/A |
| Arp5DBD+LLD | This study | N/A |
| Arp5R482A/R488A/R496 | This study | N/A |
| Arp5R514A/R516A | This study | N/A |
| Arp5R301A/R307A/R316A | This study | N/A |
| D516-564 | This study | N/A |
| Arp5.HA | This study | N/A |
| **Software and algorithms** | | |
| Alpha2 Fold | EMBL-EBI(3) | https://alphafold.ebi.ac.uk/entry/P53946 |
| PyMOL Molecular Graphics System 2.5.4 | Schrödinger, LLC | https://pymol.org |
| GraphPad Prism V.10 | GraphPad Software Inc RStudio | https://www.graphpad.com/scientific-software/prism/ |
